# Predicting the impact of sequence motifs on gene regulation using single-cell data

**DOI:** 10.1101/2020.11.26.400218

**Authors:** Jacob Hepkema, Nicholas Keone Lee, Benjamin J. Stewart, Siwat Ruangroengkulrith, Varodom Charoensawan, Menna R. Clatworthy, Martin Hemberg

## Abstract

**Background:** Binding of transcription factors (TFs) at proximal promoters and distal enhancers is central to gene regulation. Yet, identification of TF binding sites, also known as regulatory motifs, and quantification of their impact on gene expression remains challenging.

**Results:** Here we infer putative regulatory motifs along with their cell type-specific importance using a convolutional neural network trained on single-cell data. Comparison of the importance score to expression levels across cells allows us to identify the TFs most likely to be binding at a given motif. Using multiple mouse tissues we obtain a model with cell type resolution which explains 29% of the variance in gene expression. Finally, by applying scover to distal enhancers identified using scATAC-seq from the mouse cerebral cortex we characterize changes in distal regulatory motifs during development.

**Conclusions:** It is possible to identify regulatory motifs as well as their importance from single-cell data using a neural network model where all of the parameters and outputs are easily interpretable to the user.

## Background

One of the central goals in genomics is to understand how different phenotypes are determined by the DNA sequence. In higher eukaryotes the vast majority of bases represent non-coding DNA and our ability to predict the function of these sequences is incomplete. An important role for non-coding DNA is to regulate the expression of protein coding genes, and a key mechanism is through binding of TFs to proximal promoters and distal enhancers. Although the underlying principles and mechanisms of TF binding have been studied extensively, determining which motifs are functional and quantifying their importance remains a major challenge (1–3).

Identifying regulatory motifs from sequence alone is hard because in higher eukaryotes most motifs are degenerate and the number of matches in the genome is typically much larger than the number of sites bound by a TF (4). Using epigenetic information, such as TF expression levels, TF binding, chromatin accessibility, and histone modifications, it is possible to obtain more accurate models of which motifs are relevant for expression (5,6).

Today, assays for quantifying the epigenome can be carried out for single cells (7–9), resulting in datasets with finer cellular resolution. However, there is a shortage of computational methods that can take full advantage of these data to infer regulatory motifs and their impact on molecular phenotypes, such as expression levels or open chromatin.

Although there are methods for inferring regulatory motifs from single-cell data, e.g. SCENIC (10), HOMER (11) and Basset (12), none of these methods is designed for both *de novo* motif identification and quantification of the contribution to gene expression levels with scRNAseq. A variety of machine learning approaches have been applied to bulk RNA-seq and ChIP-seq data to identify sequence and epigenetic features that determine gene expression (6,13–16). In recent years, deep learning methods have become very popular (17,18) and they have been used for the task of predicting gene expression from sequence (6,19,20).

## Results

Here, we perform *de novo* discovery of regulatory motifs and their cell lineage specific impact on gene expression and chromatin accessibility from single cell data using scover, a convolutional neural network (**Fig 1a, b**). Scover takes as input a set of sequences, e.g. promoters or distal enhancers, along with measurements of their activity, e.g. expression levels of the associated genes or accessibility levels of the enhancers. The output is a set of motifs associated with the convolutional filters along with a vector of influence scores for each motif in each output cell. Scover was implemented using the PyTorch framework (21) to be compatible with the scanpy workflow (22), and it is available under the MIT license at https://github.com/jacobhepkema/scover.

**Figure 1:**
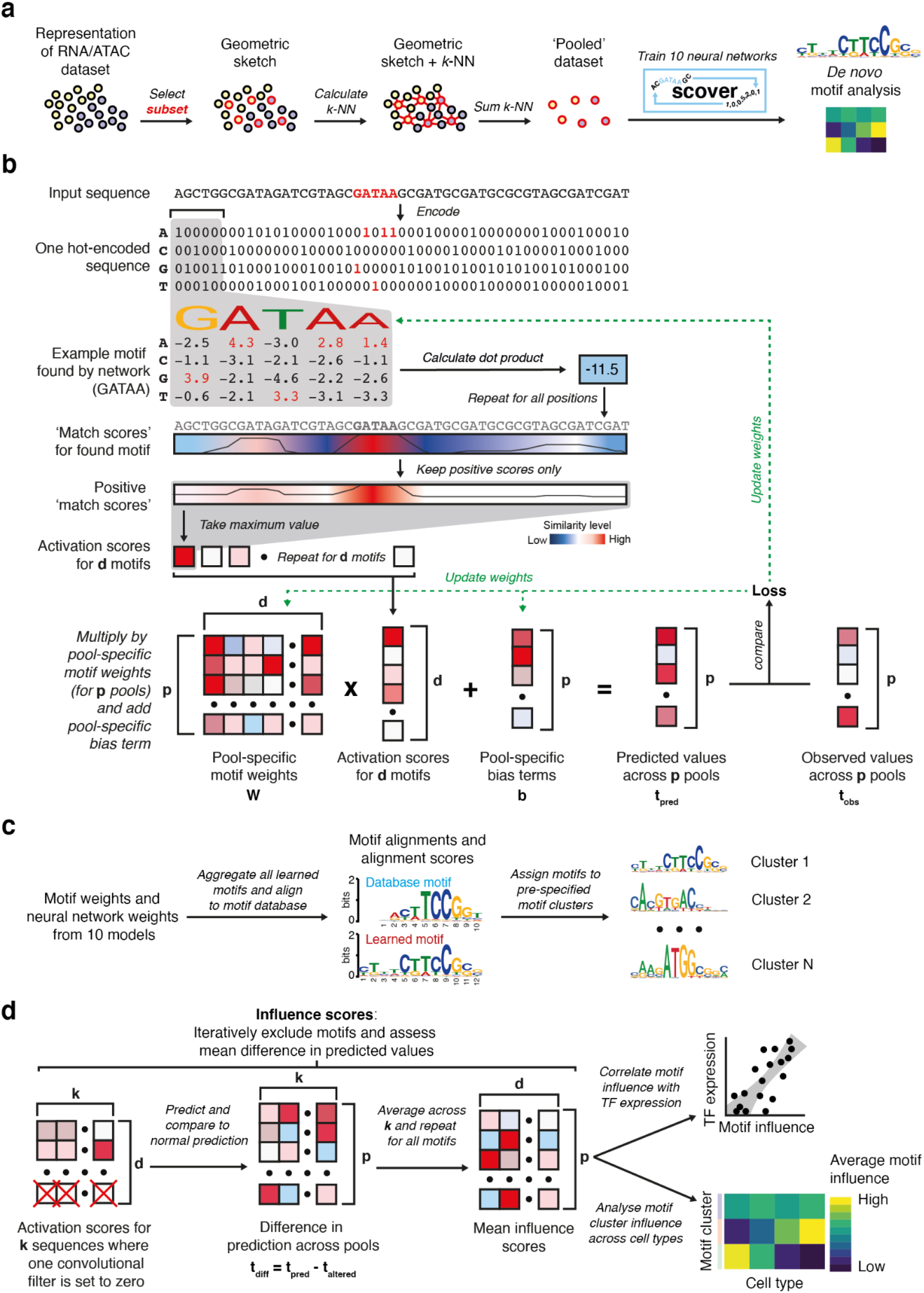
Overview of the scover workflow. (*a*) Cell pooling strategy. A subset of cells is selected using geometric sketching. The *k*-nearest neighbors for cells in the subset are summed to produce a less sparse representation of the original dataset. (*b*) Scover infers regulatory motifs that are predictive of the signal associated with a set of sequences using a neural network consisting of a single convolutional layer, a rectified linear unit, global max pooling, and a linear layer with bias term. (*c*) The identified motifs are merged and compared to annotated motifs and assigned to pre-specified motif clusters based on their most significant alignment. (*d*) To evaluate the contribution of each motif an influence score is calculated using a leave-one-out strategy.

Single-cell measurements from sequencing experiments typically have a large fraction of zeros. To overcome the accompanying challenges (23), scover reduces the sparsity by summing the values for *k*-nearest neighbors (default *k* = 100) of a set of seed cells to generate a ‘pooled’ dataset (**Fig 1a**). This strategy does not rely on existing cell type annotation, and it also retains intra-cell type variability, as opposed to taking the average for existing cell annotations. The initial seed cells are selected by geometric sketching, which evenly samples cells across a representation of the dataset, preserving rare cell states (24). The use of the *k*-nearest neighbor graph ensures that most cells in a pool will come from the same cell type, however, many pools will be heterogeneous. The cell type annotation for a pool is determined by the most abundant cell type within the pool.

We use several metrics to evaluate the effect of different pool sizes (**Fig S1**). Reassuringly, we have found that results for the datasets considered in this manuscript are robust with respect to the pool size, and it is up to the user to strike the balance between model accuracy and cell type resolution. Sequences are fed into a single convolutional layer where *d* (default *d* = 600) filters representing regulatory motifs are combined before being fed to a neural network layer resulting in a predicted expression value. The shallow architecture requires fewer parameters which allows smaller input datasets to be used and it makes it more likely that a whole motif representation will be found (25).

Since the neural network is optimized using a stochastic approach, reproducibility can be achieved by running it *r* times (default *r* = 10) and excluding motifs that are found in fewer than 50% of models (26). To facilitate the overview of the found motifs, scover automatically compares the motifs to those annotated in CIS-BP (27) (if available). Randomly initialized convolutional filters may converge to recognise similar sequences that are enriched in the sequence input. To organize the *rd* motifs, they are assigned, based on their most significant motif alignment, to pre-specified motif clusters published elsewhere (28) (**Fig 1c**).

Analysis of the convolutional filters and their associated weights can provide insights about the contribution of regulatory motifs to the observed measurements. To evaluate the importance of each motif, we employed a strategy inspired by Maslova et al (26) whereby an influence score *n*_*ip*_ is calculated for each motif *i* and pool *p* by excluding the motif from the analysis and comparing the prediction of the modified model to the original model (**Fig 1d**). Influence scores in pools corresponding to the same cell type are averaged and influence scores for motifs belonging to the same motif cluster are summed. Since influence scores tend to be largely similar across cell types, we present the *z*-transformed values to highlight cell type specific patterns.

When multiple TFs recognize the motif of a motif cluster, additional information is required to identify the most likely protein. Although transcriptome data alone is insufficient, we can rank the candidates by calculating the Spearman correlation *r*_*ct*_ between the summed influence scores for motifs in motif cluster *c* and the expression levels of TF *t* across all pools. We assume that if |*r*_*ct*_| is high, then the TF is more likely to bind to its cognate site.

### Scover identifies regulatory motifs in human kidney

We applied scover to a scRNAseq dataset from human fetal and adult kidney containing a total of 67,471 cells (9). The 1kb input sequences were 500 nts upstream and downstream of the transcription start site (TSS) of each expressed gene. We used *d =* 600, and of the 6,000 convolutional filters, 8% could be grouped into 11 reproducible families corresponding to known motifs (**Fig. 2a,b, S2**). Analysis of the top non-aligned motifs shows that many correspond to ETS motifs or GC-rich motifs, suggesting that they are likely biologically meaningful nonetheless (**Fig. S3**). Based on the validation set, the scover model explains 15% of the variance in gene expression, and the performance drops to 9% if non-matching motifs are excluded. To simplify the presentation of the results we grouped the 60 cell types into five categories: endothelial, immune, nephron epithelial, nephron progenitor and stromal (**Table S1**).

**Figure 2:**
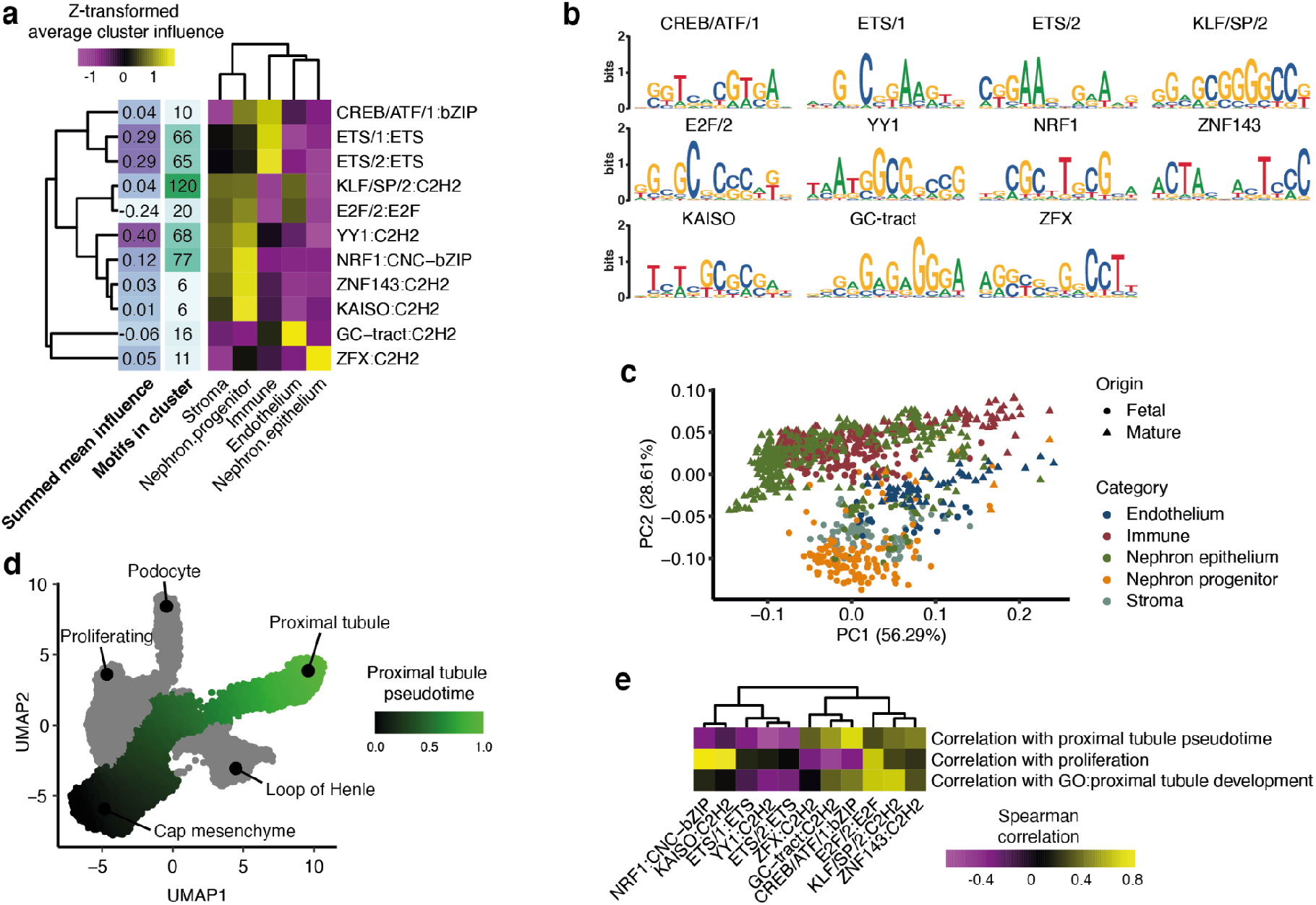
Analysis of proximal promoters for the fetal and adult human kidney. (*a*) Z-transformed influence scores for eleven motif clusters and five cell categories. (*b*) Example motif logos from the eleven motif clusters. (*c*) Projection of the pools onto the first two principal components of the influence score matrix reveals similar regulatory profiles. (*d*) Pseudotime trajectory for nephron progenitor development across single cells in fetal kidney represented as a UMAP plot. (*e*) Spearman correlations between motif influence scores and mean pseudotime values or expression of genes related to proliferation and proximal tubule development.

Dimensionality reduction using principal component analysis (PCA) reveals that the biggest difference in influence correlated with the E2F/2:E2F weights (**Fig. 2c, S4a,b**; Spearman R = 0.87), suggesting that the first principal component might be related to the cell cycle (29). The second strongest trend separates two groups, the first consisting of nephron progenitors and immune cells, and the second consisting of nephron progenitors and stromal cells (**Fig 2c**). The third principal component separates immune cells and epithelial cells from the rest (**Fig S4c**).

Since one of the largest differences in terms of influence scores is between nephron progenitors and nephron epithelium cells, we investigated the developmental trajectory by carrying out a pseudotime analysis. This revealed how the progenitors branch towards three distinct fates: podocytes, loop of Henle, and proximal tubules (**Fig 2d**). Focusing on the latter, we calculated the correlation between the expression levels of six markers associated with tubule development and the influence scores to reveal strong associations for CREB/ATF and GC-tract. This suggests that we can relate influence scores not just to discrete categories but also to continuous processes.

Although promoters share some distinct sequence features, there is also a considerable diversity to allow for differential regulation. To visualize this diversity we applied UMAP dimensionality reduction to the matrix containing the inferred motif occurrences in each promoter, and then we assigned color to each dot based on its motif score (**Fig 3a**, Methods). Interestingly, promoters separated in terms of their motif contents; KLF/SP/2 motifs were abundant across almost all promoters, some promoters were enriched for ETS/1 motifs or YY1 motifs, and promoters with high scores for ZNF143 separated from the other promoters. Similarly, by calculating the correlation between motif family influence scores across pools, we observe that the motif families fall into two major groups with GC-rich KLF and E2F motifs in one cluster, further reinforcing the notion that there is a higher order organization of motifs (**Fig S4d**).

**Figure 3:**
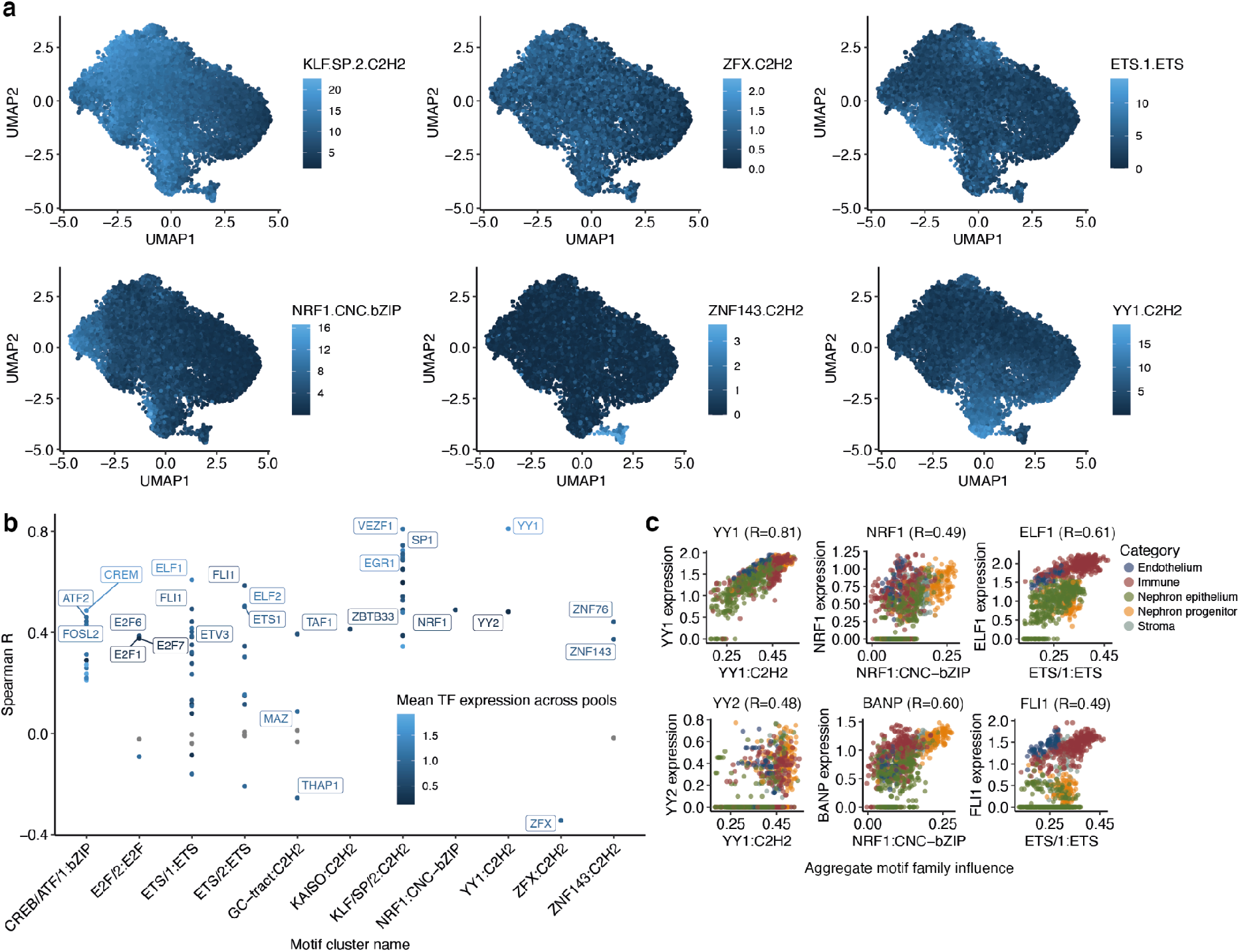
Motif cluster impact for the human kidney dataset. (*a*) UMAP plots of 18,150 promoters based on max-pooled rectified motif occurrences. Subplots show the summed max-pooled rectified motif scores for selected motif clusters in each promoter. (*b*) Spearman correlation between motif influence scores and expression levels of TFs binding to the motif in question. TFs with *p*-value>=0.05 (after Benjamini-Hochberg correction) are in gray, the significant ones are colored by expression level, and selected top TFs for each cluster are named. (*c*) Correlation between motif influence and expression of selected TFs across pools. The Spearman correlation is reported at the top of each panel.

Since multiple TFs can compete to bind similar motifs (30), we set out a strategy to identify which TFs could putatively underlie the observed influence scores for each motif family. We do this by correlating motif family influence scores with the expression levels from TFs in those motif families, under the assumption that TF expression relates to its activity.

Comparison of TF expression and influence scores reveals that 17 of the 80 TFs associated with the 11 motif families are strongly correlated with |*r*|>0.5 and a significant p-value (**Fig 3b**). These correlations can be used to pinpoint which putative TFs more likely correspond to the network predictions. For instance, while *YY1* and *YY2* have very similar motifs with an ATGG core (31), *YY1* has a higher correlation, suggesting that it is less likely that *YY2* corresponds to the network prediction (**Fig 3c**). Similarly, while *NRF1* has a moderate correlation with influence scores, it more likely corresponds to the recently characterized TF BANP that recognizes a similar CGCG motif (32), as it has a higher correlation with influence scores. However, we cannot rule out that multiple TFs correspond to the same motif cluster. For instance, *ELF1* and *FLI1* both correlate moderately with ETS/1 cluster network influence scores, which supports the notion that ETS family TFs can bind similar sites and act in functionally redundant ways (33).

### Characterization of regulatory motifs in 18 mouse tissues

Next, we applied scover to 18 tissues from the Tabula Muris (8), containing a total of 67 cell types and 38,080 cells. Only 7.55% of the 6,000 convolutional filters matched 13 different motif families, 7 of which were the same as for the human kidney (**Fig 4a,b, S5**). Again, we found that motifs that did not match any database motifs were very CpG-rich and often resembled known motifs such as ETS family motifs (**Fig S6**), and in their absence the fraction of variance explained is reduced from 29% to 18%.

**Figure 4:**
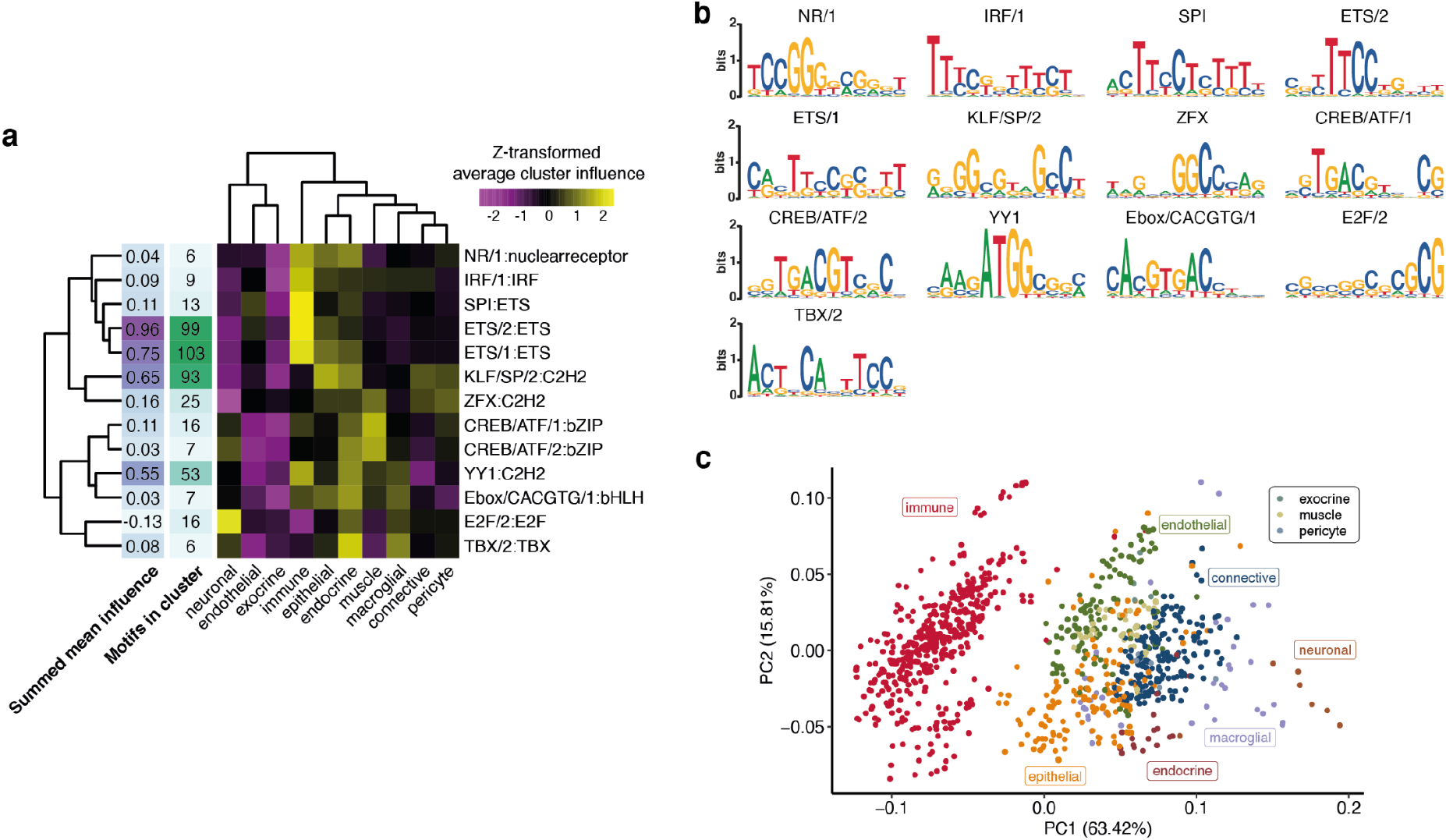
Analysis of proximal promoters from the Tabula Muris. (*a*) Z-transformed neural network influence for aggregated motif clusters. (*b*) Logos representing the thirteen motif clusters. (*c*) Pools are projected onto the space represented by the first two principal components of the motif influence matrix.

We categorized cell types as adipose, connective, endocrine, endothelial, epithelial, exocrine, immune, macroglia, muscle, neuron and pericyte (**Table S2**). Dimensionality reduction of the influence score matrix reveals three distinct groups: immune cells, neurons, and the remaining cells (**Fig 4c**), consistent with the expression analysis (34). The highest influence scores are found for ETS, followed by KLF and YY1 (**Fig S5**). We analyzed all promoters in the dataset for their motif composition (**Methods**). E2F was the most abundant motif, whereas YY1, TBX/2, and ETS/1 were specifically found in subsets of promoters (**Fig 5a**). Interestingly, there were subsets of promoters that were specifically enriched for IRF/1 and SPI only, respectively (**Fig S7**). GO-term analysis of these promoters showed an enrichment for immune-related GO terms (**Fig S7**), confirming the role of these motif families in immune cell function (35,36).

**Figure 5:**
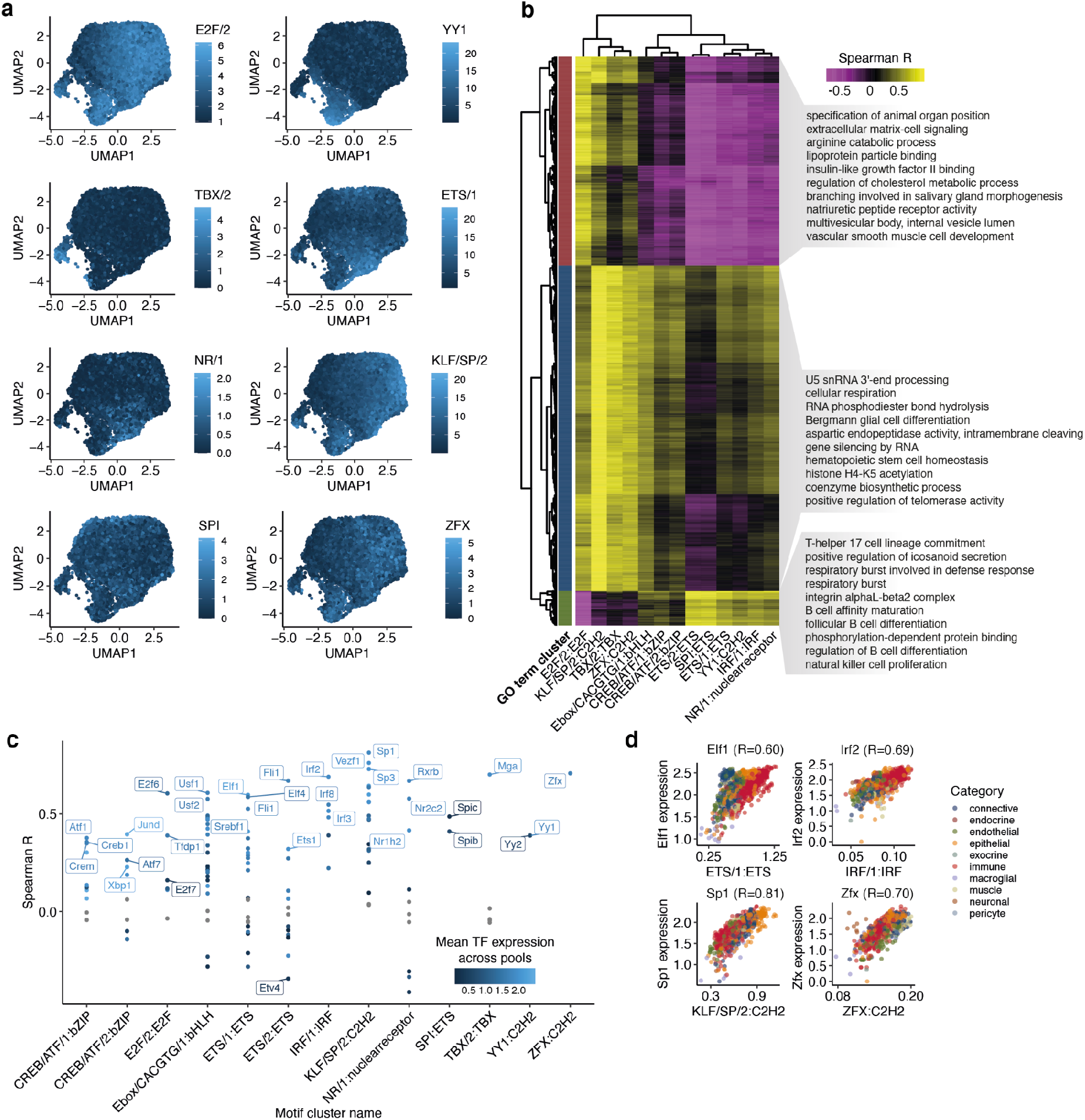
Motif cluster impact for the Tabula Muris dataset. (*a*) UMAP plots of 17,270 mouse promoters based on max-pooled rectified motif occurrences. Subplots show the summed max-pooled rectified motif scores for selected motif clusters in each promoter. (*b*) Spearman correlations between expression of GO terms and aggregate motif influences in pools. Randomly selected GO terms are shown for each cluster. (*c*) Spearman correlation between motif influence and expression levels of TFs binding to the motif in question. TFs with *p*-value>=0.05 (after Benjamini-Hochberg correction) are in gray, the significant ones are colored by expression level, and selected top TFs for each cluster are named. (*d*) Correlations between aggregate motif family influence scores in pools and expression for selected TFs. The Spearman correlation is reported at the top of each panel.

We hypothesized that in addition to having higher activity in specific cell types, the identified motifs would also be associated with distinct sets of cellular processes. We obtained 16,610 gene lists representing a diverse set of processes from the Gene Ontology (GO) database (37), and for each list we calculated the correlation between the average expression of the genes and influence scores across pools. We kept the terms with the top 1% of correlation scores, and we found that the motifs segregated into two main clusters, while the GO terms fall into three groups (**Fig 5b**). Reassuringly, the influence scores for the motifs that are highest in immune cells were positively correlated with genes associated with immune related terms. The motifs in this group (ETS, SPI, YY1, IRF) have previously been reported to be important for immune cells (13,26,35,36,38–40). The second group corresponds to GO terms that are primarily related to differentiation and morphogenesis. This group is characterized by high activity of E2F and KLF. The third group is more difficult to characterize and we are unsure about the biological interpretation.

Comparison of TF expression levels and motif activity scores reveals several known regulatory principles (**Fig. 5c**). For example, the TFs with highest correlation with ETS/1 influence scores are Elf1, Elf4 and Fli1. The high scores for ETS/1 in immune cells suggest that these TFs play a role in immune cell function. Indeed, Elf1 (41), Elf4 (42), and Fli1 (43) have all been described to play a role in regulating transcriptional programmes related to the immune system. TFs with high correlations with KLF/SP/2 influence scores include Sp1 and Sp3, which have previously been suggested to act cooperatively to regulate downstream targets (44). In other cases, expression level can distinguish between TF influences: both Yy1 and Yy2 are similarly correlated to YY1 family influence levels, and they bind similar motifs (45), but Yy1 has higher expression levels, making it more likely to be the TF with a higher influence on downstream expression.

### Identification of distal regulatory motifs in mouse cerebral cortex

To demonstrate the versatility of scover beyond analyzing promoters and scRNAseq data, we used it to infer regulatory motifs from open chromatin by analyzing data from mouse cerebral cortex collected by Chen et al using SNARE-seq (7). The cortex is organized in layers with distinct functions and we grouped cell types based on these categories. The dataset contains cells from both postnatal day 0 (P0) and adult, and we processed both time points after running the analyses separately. Since we were interested in distal enhancers, only 152,133 loci in P0 and 174,631 loci in adult that were outside the [-8 kb, 2kb] region relative to annotated TSSs were considered.

For P0 we found 19 motif clusters and for adult we found 16, with 10 found for both time points (**Fig 6a**). For P0 11% and for adult 9.65% of convolutional filters matched a known motif (**Fig S8**), and we again found that many of the non-matching filters correspond to partial matches of known motifs (**Fig S9**). During development the mouse cerebral cortex undergoes changes in gene expression (46,47). We hypothesized that there would be concomitant changes in the landscape of distant regulatory elements. Analyzing open chromatin loci across the datasets, we found that the majority of P0 and adult open chromatin regions do not overlap (**Fig S10**).

**Figure 6:**
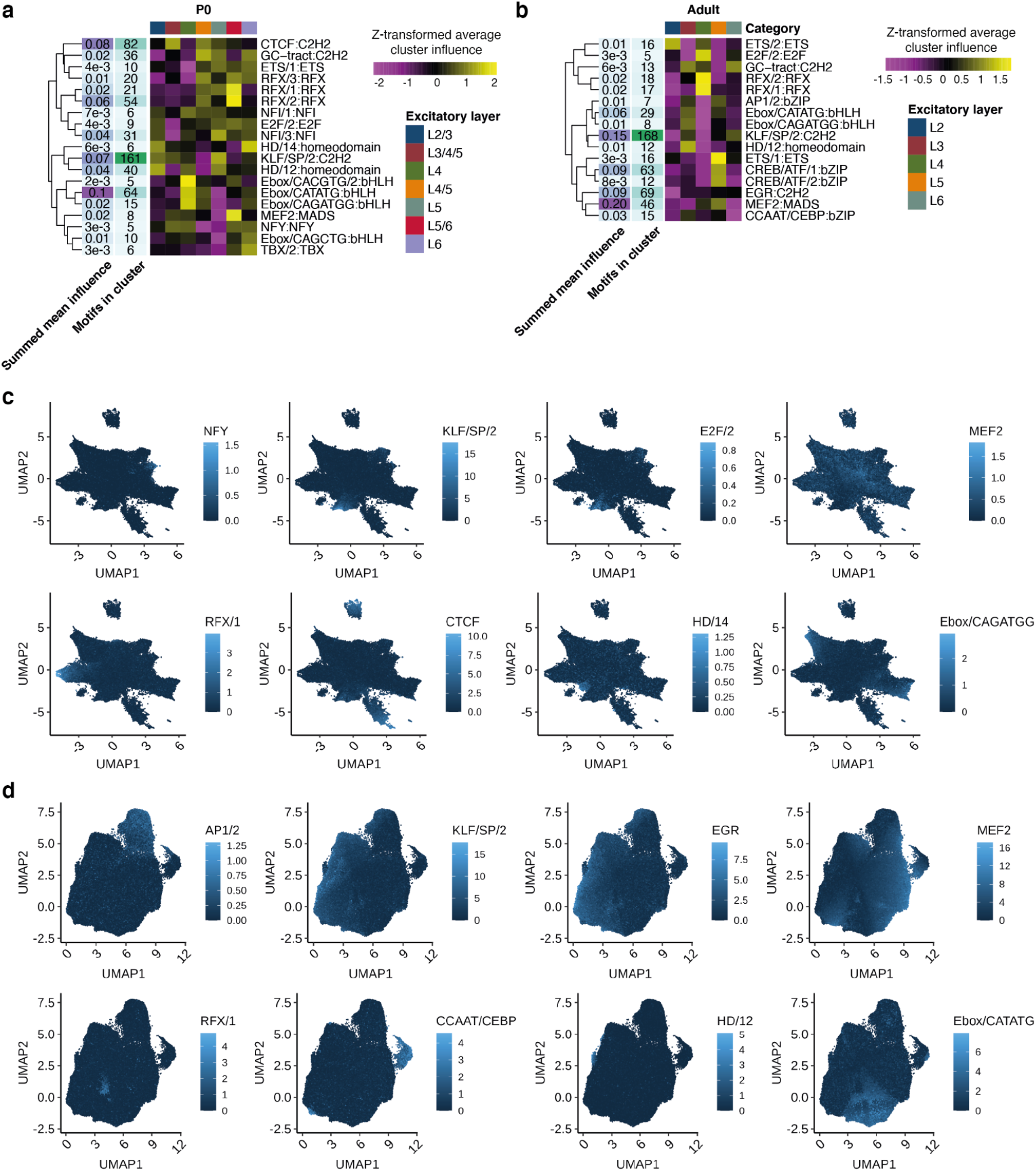
Analysis of distal open chromatin peaks from mouse cerebral cortex. (*a-b*) Z-transformed average neural network influence score per motif cluster in (*a*) P0 and (*b*) adult. (*c-d*) UMAP embeddings of regulatory regions according to the motif hits for (*c*) P0 and (*d*) adult. Color shows selected summed motif family scores in each region, except for two scores: log1p(|dist|), which shows the log-transformation of 1 plus the absolute distance to the nearest TSS, and # accessible, which shows the number of pools where the open chromatin peak is observed.

The fraction of variance explained by the model is similar to the kidney scRNAseq data with 19% for P0 and 10% for adult. As expected, the motifs identified had lower CpG-content than the ones found at promoters (**Fig S11**). We found more motifs than for the two scRNAseq datasets, suggesting a greater diversity than for promoters although it is worth keeping in mind that there were almost an order of magnitude more distal regulatory elements here. Similar to the scRNAseq data, there is a range of almost two orders of magnitude in the influence scores of the different clusters, with Ebox, CTCF, and KLF near the high end, and ETS and E2F near the low end (**Fig S12**).

Comparing the P0 and adult datasets, we identified several differences in found motif families. For instance, CREB and EGR were among the identified motif families in adult, but not in P0 mice. TFs from these families have been implicated in neuronal plasticity (48–51), and Egr1 expression goes up drastically in neurons throughout cortical development (52). Similarly, CCAAT/CEBP motifs were found in the adult, and one of its associated TFs is Hlf which increases in expression during mouse development and is implicated in brain development (53). Taken together, the analysis reveals several differences in terms of distal regulatory motifs between P0 and adult mouse cerebral cortex, but more work and data is required to validate and interpret these differences.

To identify groups of enhancers with similar motif composition, we embedded the rectified motif scores for each region using UMAP (**Fig 6c, d**). There were large differences in how widely a motif family was observed across peaks: MEF2 motifs were spread across a large number of loci in both P0 and adult mouse peaks, and EGR was widely observed in adult mouse peaks. Conversely, hox transcription factor motifs from HD/12 and HD/14 motif families had a more localized pattern, suggesting that they activate smaller subsets of loci with distinct regulatory motifs. Interestingly, one group of regions constitute a clear group of outliers in the UMAP for P0 mouse peaks and we find that they are characterized by high scores for the CTCF-motif (**Fig 6c**). This result suggests that insulator regions differ in terms of their sequence composition compared to other distal open chromatin regions.

## Discussion

Here, we have presented a convolutional neural network model, and an exploration of inferred regulatory motifs and their activities across cell types using single cell data. Previous attempts to develop quantitative models of regulatory motifs have been restricted to tissue level or cell lines (6,13), or to immune cells (26,38), and they frequently use histone marks rather than sequence to predict expression levels (54). Unlike existing motif analysis methods which mainly identify overrepresented motifs in sequences, scover also determines the relative impact of each motif on gene expression. By comparing to gene expression levels these scores can be used to pinpoint putative TFs that are related to the observed pattern.

Since we used a *de novo* strategy for finding motifs we expect that it will be especially helpful for researchers studying organisms where regulatory motifs are poorly annotated. For mouse and human non-matching motifs make a meaningful contribution, but from our results it is not yet clear what they represent. Some of them are likely good matches that have failed to converge, whereas others pick out CpG dinucleotides which are known to be associated with higher expression at promoters (6). Our results are also consistent with other studies that have categorized TFs as either preferentially binding promoters or enhancers (38). One such example is HNF4A, which is important in the kidney (55), but most likely due to its preference to distal enhancers it was not identified by our method (56).

Interestingly, there are several trends that are common to the kidney and the Tabula Muris datasets. Immune cells have regulatory profiles distinct from other cell types and they have the highest influence for ETS motifs, while KLF and ZFX have low impact (**Fig 2a, 4a**). More generally, for the seven motif clusters (CREB/ATF/1, ETS 1 & 2, KLF, E2F, YY1, ZFX) that are found in both datasets, the influence scores are in good agreement with the highest impact for the combined ETS families, followed by YY1 or KLF. A notable difference is that the quality of the fit provided by the Tabula Muris is substantially better. Since the two datasets have similar number of cells and similar sparsity after pooling, we conjecture that this is due to the greater diversity of cell types in the Tabula Muris and the use of different experimental platforms.

Although larger and more deeply sequenced datasets could allow for the discovery of additional motif clusters and better model fits, there are inherent limitations to the accuracy by which gene expression can be predicted from endogenous sequences. First, studies of gene expression noise have estimated that some of the variability is inevitable due to random fluctuations (57). Second, there are many other mechanisms of gene regulation that are not included in proximal sequence data, e.g. transcript sequence features that influence mRNA stability (58), TF concentrations, distal enhancers, histones, miRNAs, and RNA binding proteins. Third, we have only used one promoter for each gene, even though it is well known that many genes have alternative TSSs which could drastically impact regulation. Nevertheless, our results are consistent with a recent study of mouse hematopoietic cells which used epigenetic data to estimate that ∼50% of the variance in expression can be explained by the endogenous promoter (54).

## Conclusions

We have demonstrated that a convolutional neural network can be used to simultaneously infer regulatory motif sequences and their relative importance from single-cell data. Using scRNAseq data, this allows us to find TF binding motifs at promoters along with their impact on gene expression levels. Application to data from human kidney and multiple mouse tissues revealed common regulatory patterns. Finally, we demonstrated that the framework can also be used for scATACseq data, and analyses of data from the mouse visual cortex allowed us to characterize motifs found at proximal and distal regulatory regions across development.

## Methods

### Pooling

To reduce the sparsity, pools are created by grouping K-nearest neighbors of an initial selection of cells (‘seed cells’) calculated using a shared PCA embedding into *p* equally sized groups of *q* cells (**Fig 1a**). The expression values are the log-transformed sum of the non-log-transformed expression values of the single cells in that pool. The ‘seed cells’ for the pooled dataset are selected using geometric sketching (24), which evenly samples cells from an embedding of the single cell dataset. The embedding selected for ‘seed cell’ sampling is slightly different for the RNA and ATAC datasets. As the RNA datasets consist of concatenated datasets of different batches, batch-balanced K-nearest neighbor algorithm BBKNN (59) was run with default parameters, which was used subsequently to calculate a batch-balanced UMAP dimensionality reduction. 1000 ‘seed cells’ were sampled using geometric sketching for both the human kidney and Tabula Muris datasets across this batch-balanced UMAP. For both P0 and adult SNARE datasets, 100 ‘seed cells’ were sampled for each dataset using a PCA embedding.

### One-hot encoding of nucleotide sequences

For the pooled transcriptomic datasets the sequences correspond to 500 bp upstream and 500 bp downstream of gene TSS (for a total size of 1000 bp per sequence). For the Tabula Muris and SNARE-seq datasets, sequences were isolated from the mm10 reference genome using the Ensembl v.79 annotations (60). For the human kidney dataset, sequences were isolated from the hg38 reference genome using the Ensembl v.86 annotations. For the pooled accessibility datasets, each sequence corresponded to 250 bp around the center of the peak region. Input nucleotide sequences are one-hot encoded into a 2D matrix such that the columns correspond to [A, C, G, T], respectively. For each position in the sequence, the corresponding row in the matrix will have a 1 in the column corresponding to the original nucleotide and a 0 otherwise (**Fig. 1a**).

### Convolutional neural network architecture

Scover is a convolutional neural network composed of a convolutional layer, a rectified linear unit (ReLU) activation layer, a global maximum pooling layer, and a fully connected layer with bias and multiple output channels. The convolutional layer takes as input the one-hot encoded DNA sequences, and the fully connected layer outputs predictions across the pools. Sequentially, the neural network carries out the following operations:

1. 2D convolution without bias values with *d* convolutional filters of size (*m*, 4)
2. A ReLU activation layer that sets negative values to 0
3. A global maximum pooling layer that takes the maximum value for each of *d* rectified convolutional operations
4. A fully connected layer with bias values with *d* inputs and *p* outputs

The number of output pools *p* depends on the number of cells per pool *q*. The default values for running the model are *d* = 600, *m* = 12 and *q* = 100. However, it is recommended that the user explore different combinations.

### Training the neural network

The neural network is trained in two stages: Bayesian hyperparameter optimization, and a training stage using the best hyperparameters.

In the first stage, the hyperparameters are optimized through Bayesian hyperparameter optimization (61,62). The hyperparameter search is implemented using Ray Tune (63) and hyperopt (64) and runs distributed across available GPUs. For hyperparameter tuning, we use the ASHA algorithm, which applies aggressive early-stopping to trials that do not seem to produce good results (65), finding optimal hyperparameters faster than random search.

The division of the dataset into training, test, and validation sets was performed using cross-validation. The dataset was divided into *K* folds (default: 10) to create the ‘outer test’ set (10% of data each) and the ‘inner’ sets (10% of data each). Each ‘inner’ set was consequently split into a training and a validation set of 80% and 20% of the ‘inner’ set respectively. The ‘inner’ sets were used for the hyperparameter search for each outer fold.

The hyperparameter search was inspired by a previous neural network (62). The neural network parameters were initialized *num_calibrations* (default: 100) times for each fold with weights suggested by hyperopt based on previous runs for the fold. The prior for the learning rate is log uniformly distributed over the interval [*epsilon_min, epsilon_max*] (default: [5e-4, 5e-2]). The prior for the convolutional filter weights is normally distributed with mean 0 and standard deviation *sigma_motifs*^2^, where *sigma_motifs* is sampled from a log uniform distribution over the interval [*sigma_motifs_min, sigma_motifs_max*] (default: [1e-7, 1e-3]). Similarly, the prior for the fully connected layer weights is normally distributed with mean 0 and standard deviation *sigma_net*^2^, where *sigma_net* is sampled from a log uniform distribution over the interval [*sigma_net_min, sigma_net_max*] (default: [1e-5, 1e-2]). The fully connected layer biases are initialized with the value 1e-5. The batch size was chosen by hyperopt from choices [64, 128, 256, 512]. Each network is trained for a number of epochs (default: 40). For each fold the lowest validation set errors are calculated. The optimal network initialization is chosen as the network initialization with the lowest mean error for the validation set.

During the training stage, the network is initialized using the optimal network initialization (the learning rate, batch size, *sigma_motifs*, and *sigma_net*) for each fold. For each fold, the network is trained for the same amount of epochs as before on the inner training set, and network parameters for each fold are saved at the time point of lowest validation set error. The network uses a mean squared error loss function, and the network parameters are optimized using the Adam optimizer (66). The network is trained with an early stopping strategy with patience 1. Prediction metrics are calculated for each fold separately using the prediction for the held-out ‘outer’ test set.

To ensure that the motifs identified by scover capture biologically meaningful signals, we carry out a control experiment where we train scover using a dataset where the nucleotide order of the input sequences is permuted. Small differences in correlation between the predicted and observed values for the real and permuted data is a strong indication that the identified motifs are spurious matches and that the model has failed to identify an informative model. For all of the datasets considered in this manuscript, the mean difference in the squared Pearson correlation using the real and permuted data was 0.098 for the kidney dataset, 0.21 for the Tabula Muris dataset, 0.120 for the p0 SNARE-seq dataset, and 0.063 for the adult SNARE-seq dataset. Of these, the smallest changes are for the SNARE-seq datasets, where the mean squared Pearson correlation went down by 63% for both datasets.

### Alignment of motifs

To extract position weight matrices corresponding to the convolutional filters, rectified activations are calculated for each model by convolving the convolutional filters across the training set sequences and passing the output through a ReLU layer. Consequently, for each sequence, the subsequence of length *m* that generates the highest positive activation is isolated for each convolutional filter. Sequences that do not generate a positive activation score for a given convolutional filter are not included. For each convolutional filter, the scores are converted to position frequency matrices (PFMs) of size 4 x *m* by summing the occurrences of each nucleotide at the corresponding sequence position. The PFMs are converted to position probability matrices (PPMs) by dividing each entry in the PFM by the sum of the nucleotide frequencies for the corresponding sequence position. The concatenated PPMs of the 10 models are then stored in a single MEME-formatted file. The motifs are aligned to MEME-formatted motif database files from CIS-BP (27) using the Tomtom tool from the MEME suite (67) using the argument *-*thresh 0.05. Sequence logos were visualized using R package *ggseqlogo (68)*.

### Leave-one-out analysis and influence scores

After training *r* models, each model is reinitialized using the parameters from the optimal training point (the point at which the MSE on the validation set was the lowest). The prediction for motif *i* in pool *j* and sequence *k* is denoted *b*_*ijk*_. We then set all parameters related to motif *i* to zero and calculate the perturbed score *c*_*ijk*_ for each pool. The influence score for motif *i* in pool *j* is calculated as 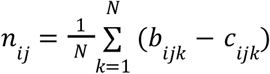 where *N* is the number of sequences in the validation set. The influence scores from *d* motifs from *K* models are concatenated to get a matrix with *dK* rows and *p* columns. For each motif, the mean influence per cell type is calculated as an average from the corresponding pools. For visualization purposes we also calculate a z-transformed version (based on cell types). Aligned motifs were assigned to a previously specified set of motif clusters (28). To ensure reproducibility, motif clusters are excluded if they contain motifs from less than half of the *K* models. Lastly, for most of the downstream analysis, motifs that did not align to known motifs are excluded. Influence scores for motifs that belong to the same motif family were summed to generate an ‘aggregate motif family influence’ matrix with *num_motif_families* rows and *p* columns. Similar to before, this matrix is then converted to include mean aggregate motif family influence scores across cell types or cell type categories, and for visualization purposes, these scores are z-transformed.

### Sequence ‘motif space’ analysis

For the kidney and tabula muris experiments, promoter analysis was carried out by iterating over all sequences using the 10 models: the calculated scores were the rectified max-pooled outputs of the convolutional layer for each model. The outputs were concatenated in the motif dimension to create an array of size (n_genes x 6000). “Motif space” UMAPs were calculated using this matrix using R package *umap* (69).

### Analysis of region overlap

The R packages ComplexHeatmap (70) and GenomicRegions (71) were used to calculate and plot the overlap in base pairs between the 250bp regions in the datasets used to train, validate, and test the models for SNARE P0 and adult.

### Proliferation and proximal tubule marker analysis

We used the following genes as markers of cell proliferation: MKI67, PLK1, E2F1, FOXM1, MCM2, MCM7, BUB1, CCNE1, CCND1, CCNB1, and TOP2A (72). The proliferation score of each pool was calculated as the average expression of this set of genes in that pool. Similarly, we used DLL1, ACAT1, PKD1, NOTCH2, AQP11 and HEYL for the proximal tubule signature genes.

### Gene Ontology analysis

The GO terms for mouse were downloaded from Ensembl Biomart. We used all categories with fewer than 50 genes for the analysis. For each GO term, the correlations between the expression of the GO term genes (across pools) and the aggregate motif influences of each motif family (across pools) were calculated. The top 500 GO terms in terms of their maximum correlation were hierarchically clustered based on euclidean distance, after which the tree was cut with *k* = 3 to form the clusters.

### Analysis of co-occurring motif families

To investigate co-occurring motif families in the Tabula Muris dataset, the max-pooled rectified outputs of the convolutional layers were first aggregated per motif family. Then, sets of genes with motif occurrence were selected for each motif family by filtering for genes with a score above half of the maximum gene score for that motif family. The upset plot was plotted using the R package UpsetR (73).

### Pseudotime analysis

The developing nephron compartment of the fetal kidney dataset was subset to cells with a developmental relationship to cap mesenchyme (excluding ureteric bud). A two dimensional UMAP embedding was calculated in scanpy (22), and was clustered using k-means clustering (k=10). Pseudotime trajectories were computed with Slingshot (74) using the cluster corresponding to cap mesenchyme as a starting cluster. Proximal tubule pseudotime values were generated for cells along a continuous trajectory path from cap mesenchyme to proximal tubule. The pseudotime values were averaged across the same cells that generated the pools in the combined human kidney dataset. In some cases, not all the cells in the pool had corresponding pseudotime values, as some cells fell outside of the k-means clusters along the principal curve corresponding to the proximal tubule pseudotime trajectory. The pseudotime values were only calculated for pools that had at least 100 cells (out of 120 cells per pool) with pseudotime values. The pooled (averaged) pseudotime values were subsequently normalized to range [0,1].

## Data processing

### Human kidney dataset

The fetal and adult kidney datasets were concatenated in the cell dimension to create a matrix of 33,660 features by 67,465 cells. A PCA was run and the first PC was ignored for further analysis as this correlated heavily with the total amount of RNA counts in the cell. The pooling strategy is described above. After pooling the datasets with *q* = 120 cells per pool, the dataset was filtered by excluding genes that were expressed in < 5% of pools. Another filter was applied: genes that were expressed in < 16% of pools were only retained if the mean non-zero log-transformed expression was higher than log10(3). This resulted in a dataset of 18,150 genes by 1,000 pools. Running a full hyperparameter sweep with cross-validation using *d* = 600 and *m* = 12 took a few hours when run across 8 NVIDIA Tesla V100 GPUs.

### Tabula Muris dataset

20 Tabula Muris FACS-sorted Smart-Seq2 datasets were concatenated in the cell dimension. 297 erythrocytes were excluded from further analysis given their low counts, resulting in a matrix of 23,433 genes by 44,652 cells. The pooling strategy is described above. After pooling the dataset with *q* = 80 cells per pool, the data was filtered by removing genes that were expressed in <3% of pools. Another filter was applied: genes that were expressed in < 8% of pools were only retained if the mean non-zero log-transformed expression was higher than log10(4). The final pooled dataset had a size of 17,270 genes by 1,000 pools. When training the model we used *d* = 600 and *m* = 12.

### SNARE-seq dataset

The P0 and adult mouse datasets were both pooled with *q* = 100 cells per pool using the strategy described above. Both datasets were filtered by excluding peaks that were present in < 6% of pools. To ensure that only distal peaks were included we removed any peak found in the [-8kb, 2kb] region relative to any TSS found in the Ensembl annotations for mm10 (v.96). Furthermore, we excluded all non-neuronal cell types from the analysis, but we included the inhibitory neurons when fitting the model even though only the excitatory neurons were used for the downstream analyses (**Table S4**). This resulted in a final pooled dataset size of 152,133 peaks by 100 pools for the P0 mouse dataset and 174,631 peaks by 100 pools for the adult mouse dataset. When training the model we used *d* = 600 and *m* = 12 for both P0 and adult mouse datasets.

## Declarations

### Ethical approval and consent to participate

Not applicable

### Availability of data and materials

### Code availability

The neural network is implemented in pytorch, examples and code are available at https://github.com/jacobhepkema/scover. Code that was used to generate the figures is available at https://github.com/jacobhepkema/scoverplots. Plots were generated using ggplot (75), pheatmap, gghalves, and other packages described above.

### Data availability

The human kidney datasets were downloaded from https://www.kidneycellatlas.org/. We downloaded 20 Tabula Muris FACS-sorted Smart-Seq2 datasets from https://doi.org/10.6084/m9.figshare.5829687.v8. We downloaded the SNARE-seq data from the Gene Expression Omnibus database under accession number GSE126074.

### Author contributions

The project was conceived by JH, NKL and MH. JH, NKL and SR wrote the code. JH and MH analyzed the data. BJS and MRC assisted with the analysis of the kidney data. VC, MRC and MH supervised the research. JH and MH wrote the manuscript with input from the other authors.

## Acknowledgements

We would like to thank Song Chen and Yingxi Lin for helpful discussions about the SNARE-seq data and Irina Abnizova, Ilias Georgakopolous-Soares, Leopold Parts and Nikos Patikas for comments and suggestions on the manuscript and the package. We are also grateful to Simon Murray for testing the repository and to Jimmy Lee for sharing code.

## Supplementary figures

**Figure S1:**
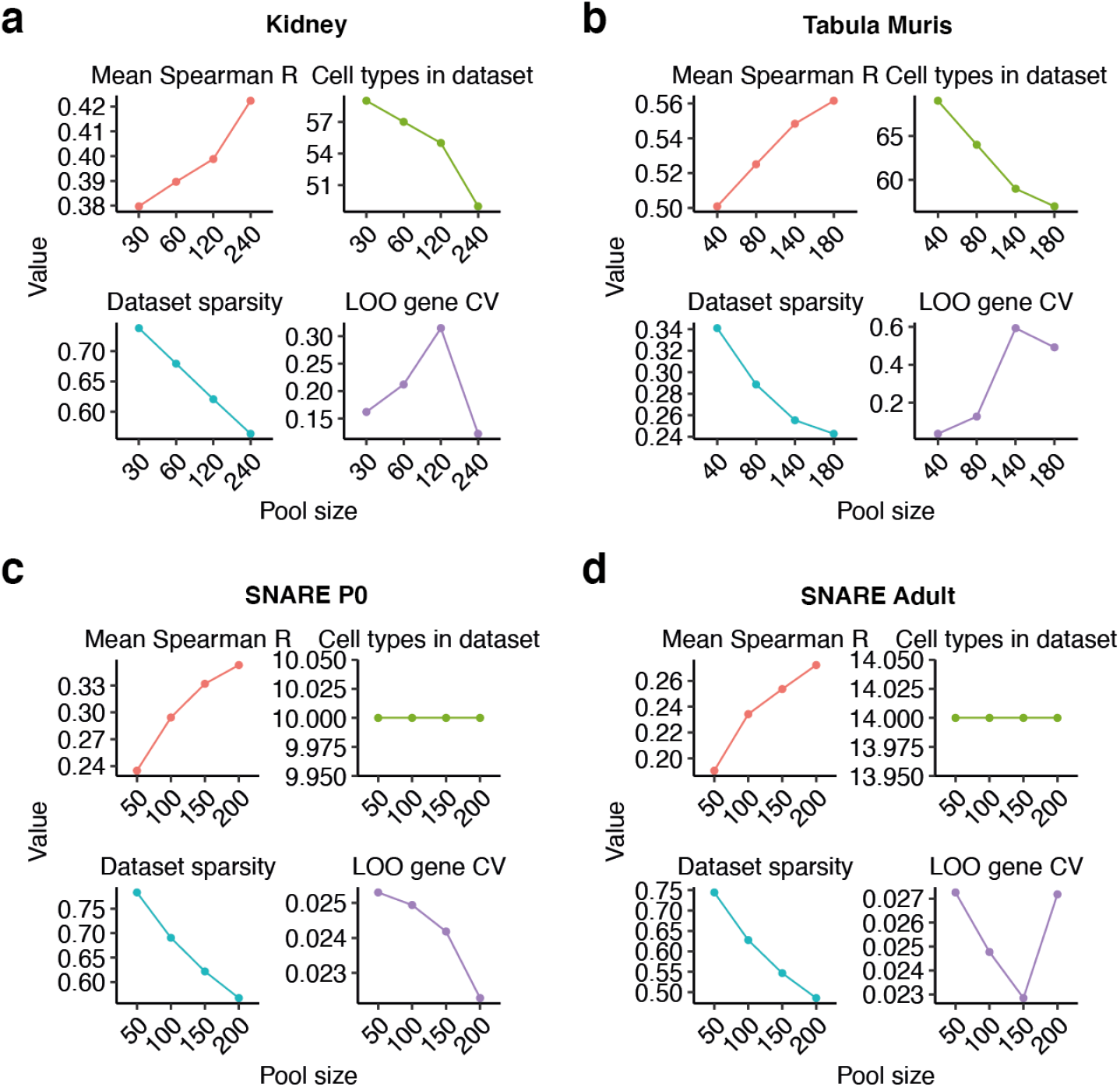
Correlations, number of cell types, sparsity (fraction of genes that are zero) and gene-wise coefficient of variation for influence scores (LOO gene CV) for different pool sizes for the datasets considered in this manuscript. The gene-wise coefficient for variation for influence scores was calculated only using reproducibly found motif families.

**Figure S2:**
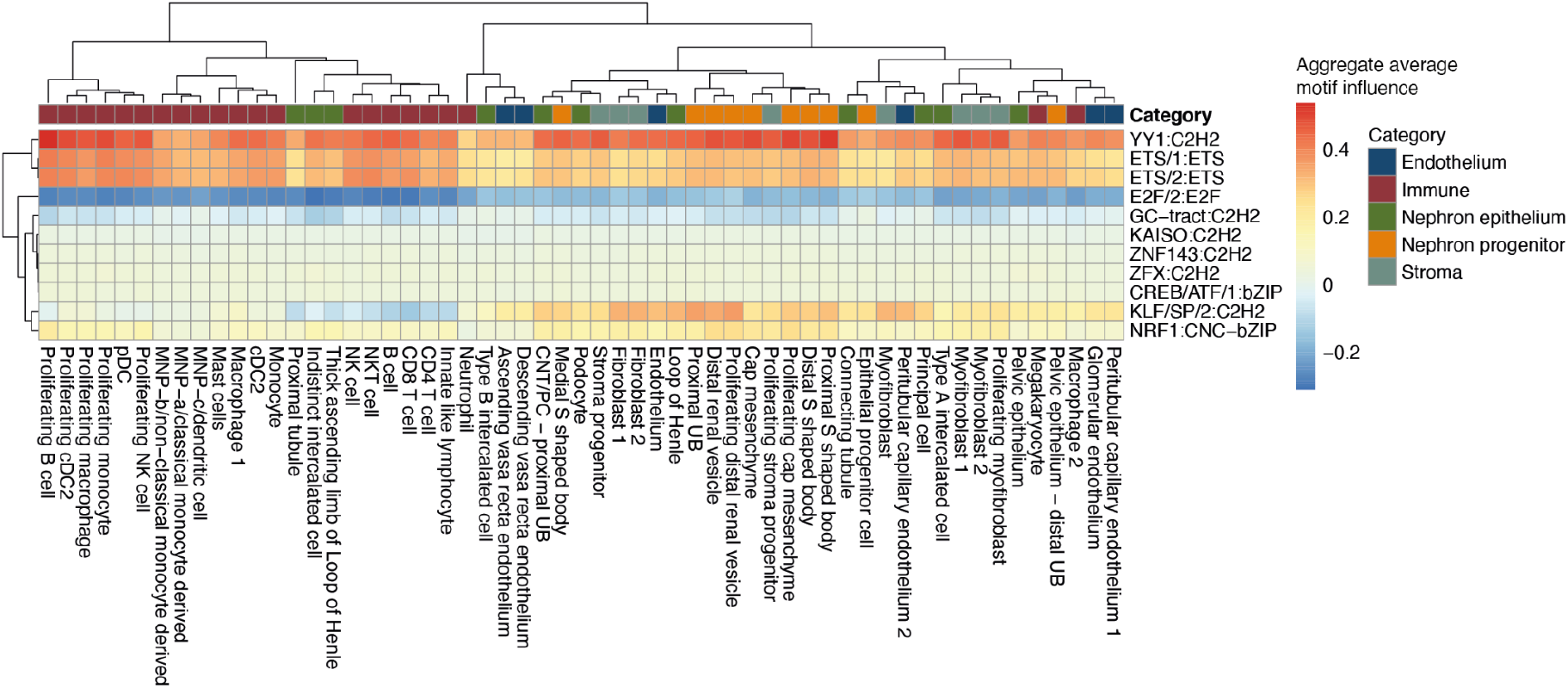
Aggregate motif weights of motif clusters averaged over cell types for the human kidney dataset.

**Figure S3:**
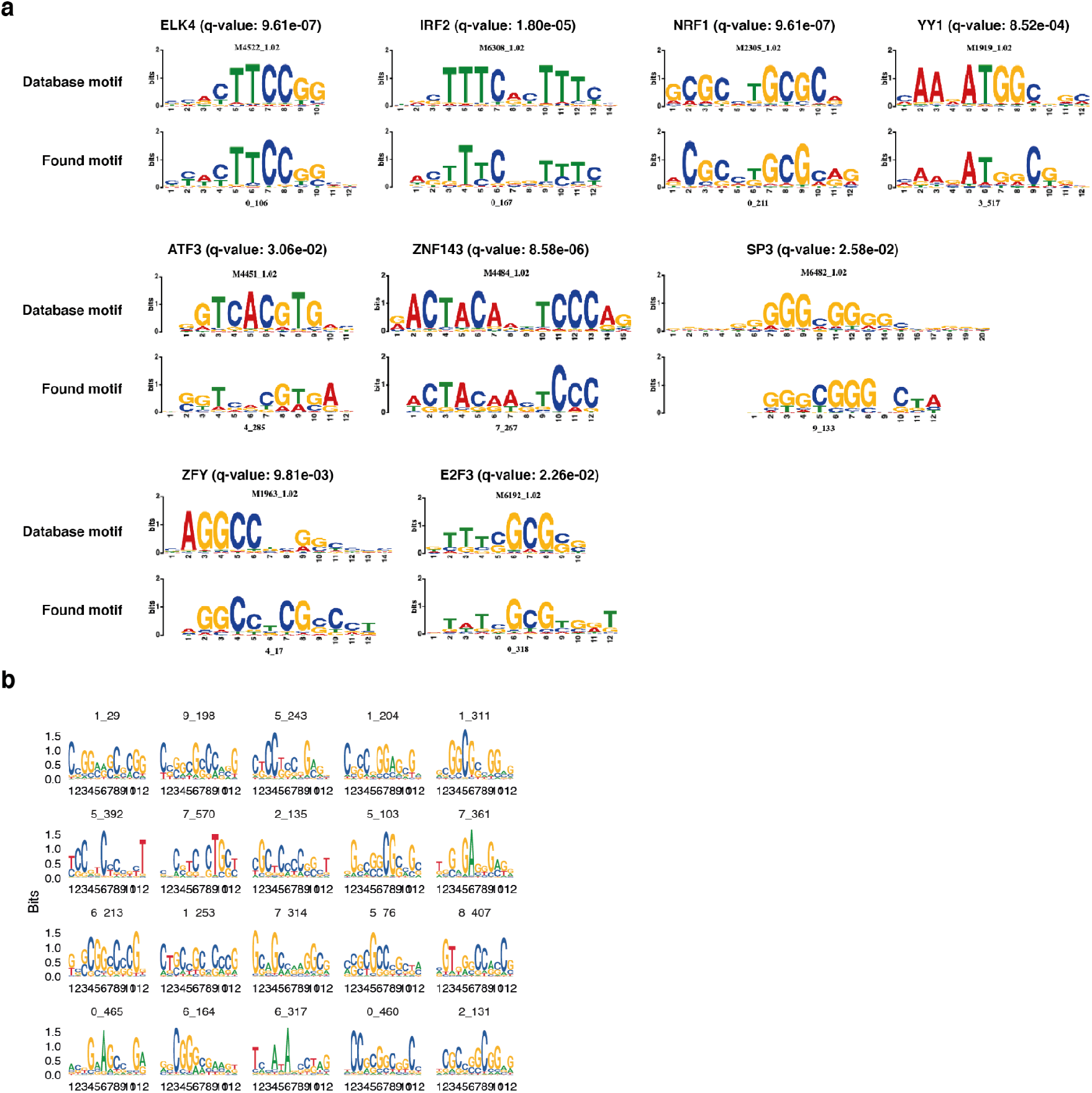
(*a*) Example motif alignments for the human kidney dataset. (*b*) Random selection of 20 motifs with high impact scores that do not significantly align to the CIS-BP database.

**Figure S4:**
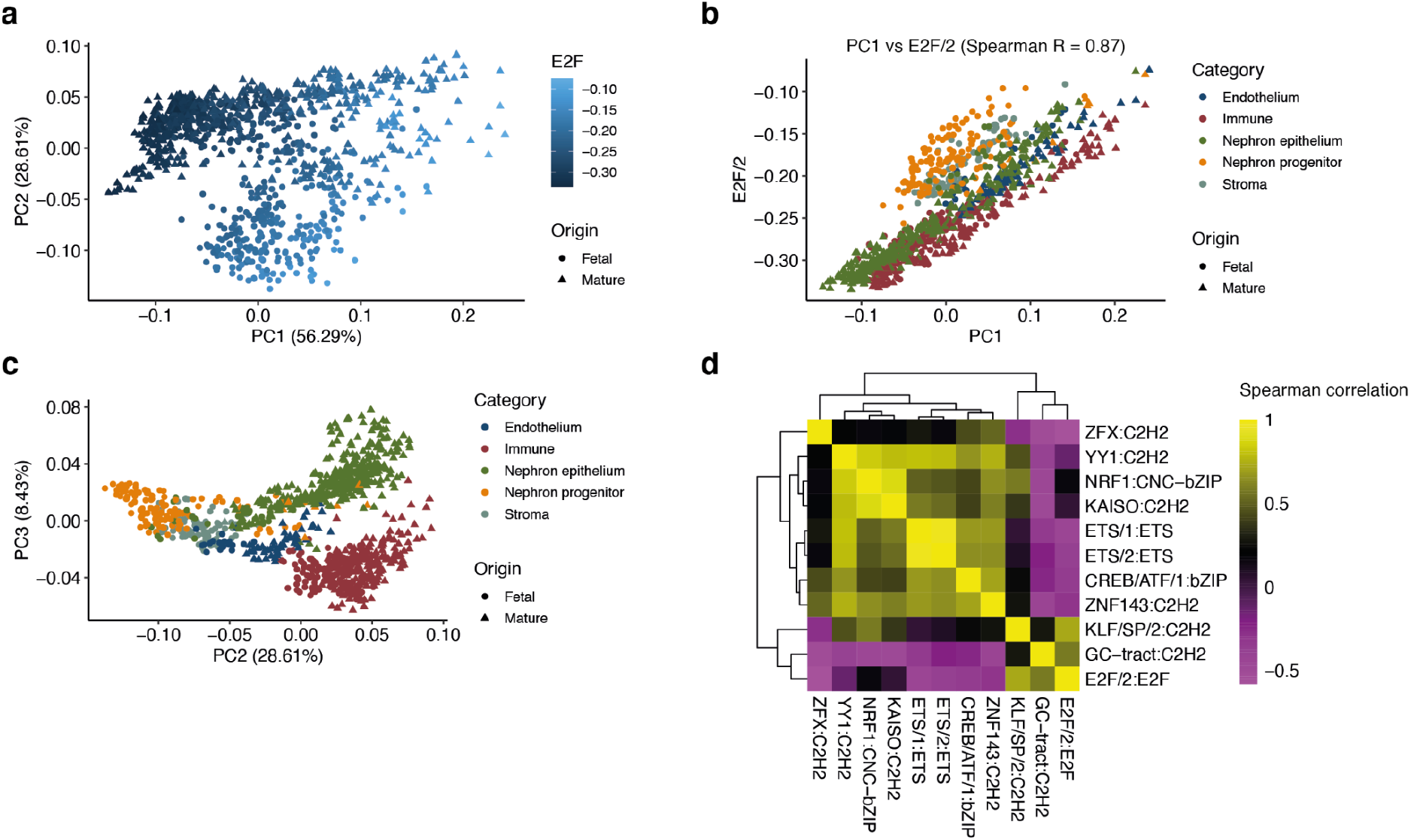
(*a*) Projection of the pools onto the space represented by the first two principal components of the influence score matrix for the human kidney dataset. Color shows E2F/2:E2F family influence scores. (*b*) Scatterplot of the first principal component of the influence score matrix and the E2F/2:E2F family weights. (*c*) Projection of the pools onto the space represented by the second and third principal components of the influence score matrix for the human kidney dataset. (*d*) Pairwise spearman correlations between motif cluster influence scores for the human kidney dataset.

**Figure S5:**
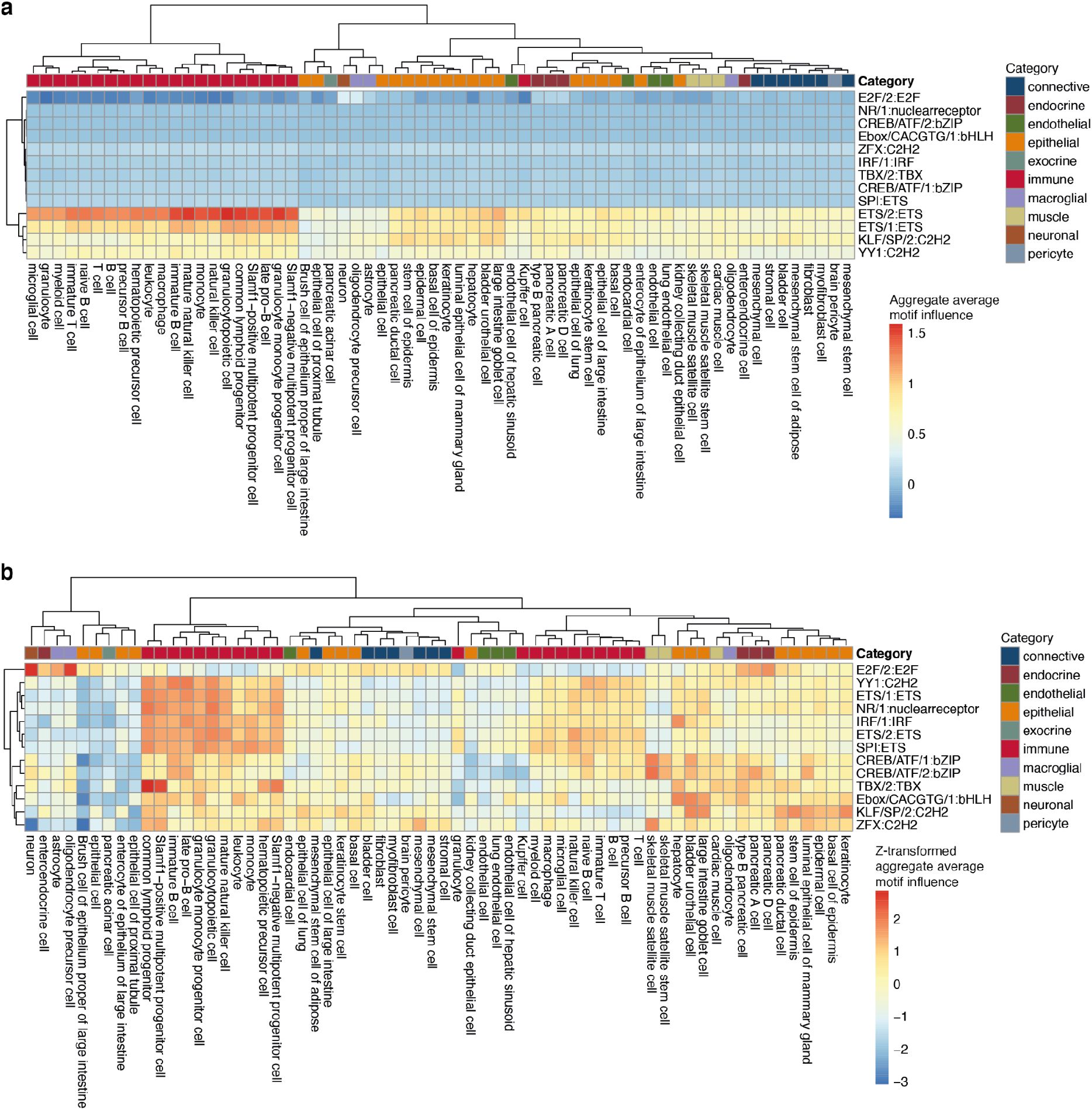
(*a*) Aggregate motif weights of motif clusters averaged in cell types for the Tabula Muris dataset. (*b*) Z-transformed aggregate motif weights of motif clusters averaged in cell types for the Tabula Muris dataset.

**Figure S6:**
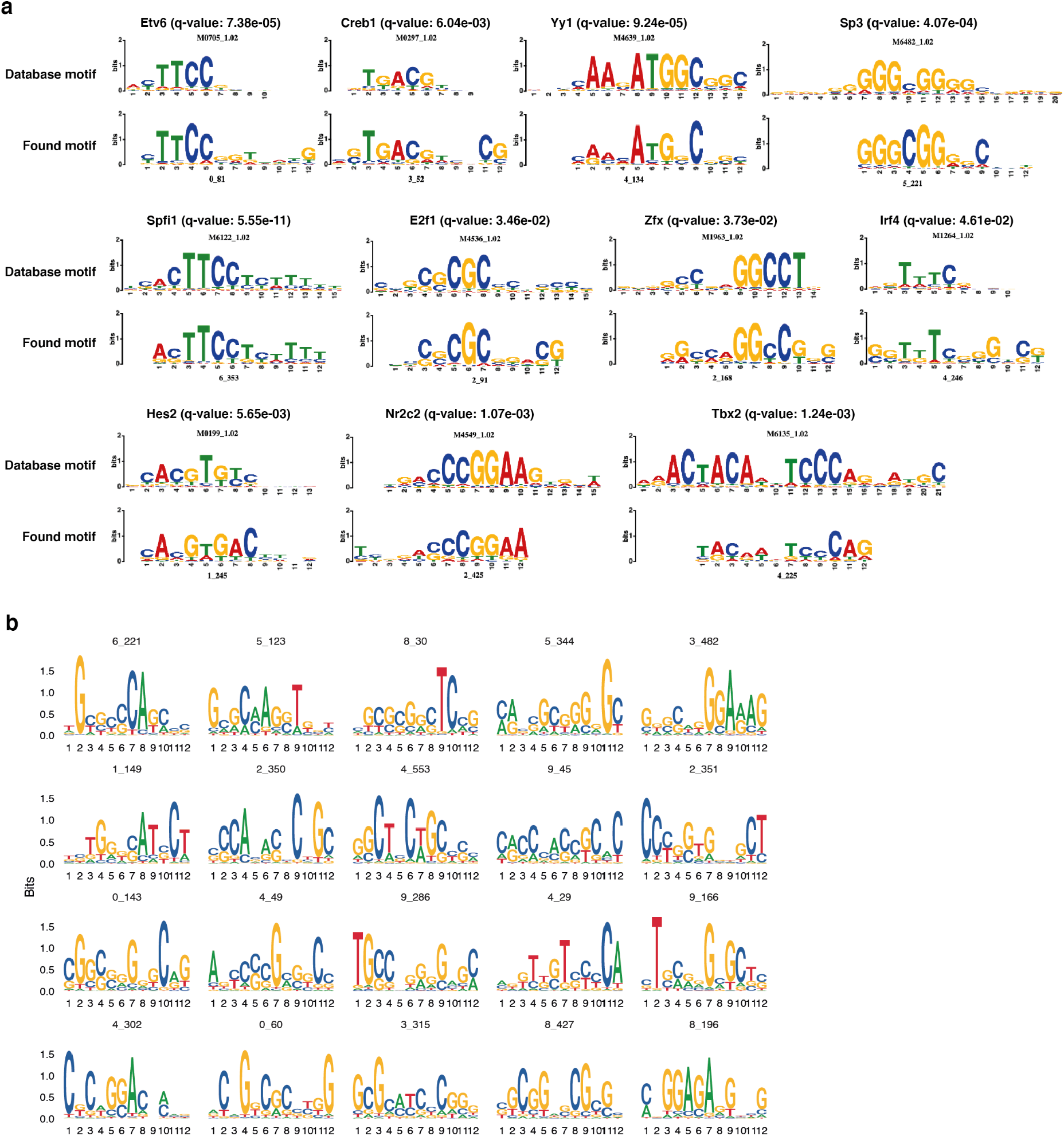
(*a*) Example motif alignments for the Tabula Muris dataset. (*b*) Examples of randomly selected motifs with high influence scores that did not align to CIS-BP.

**Figure S7:**
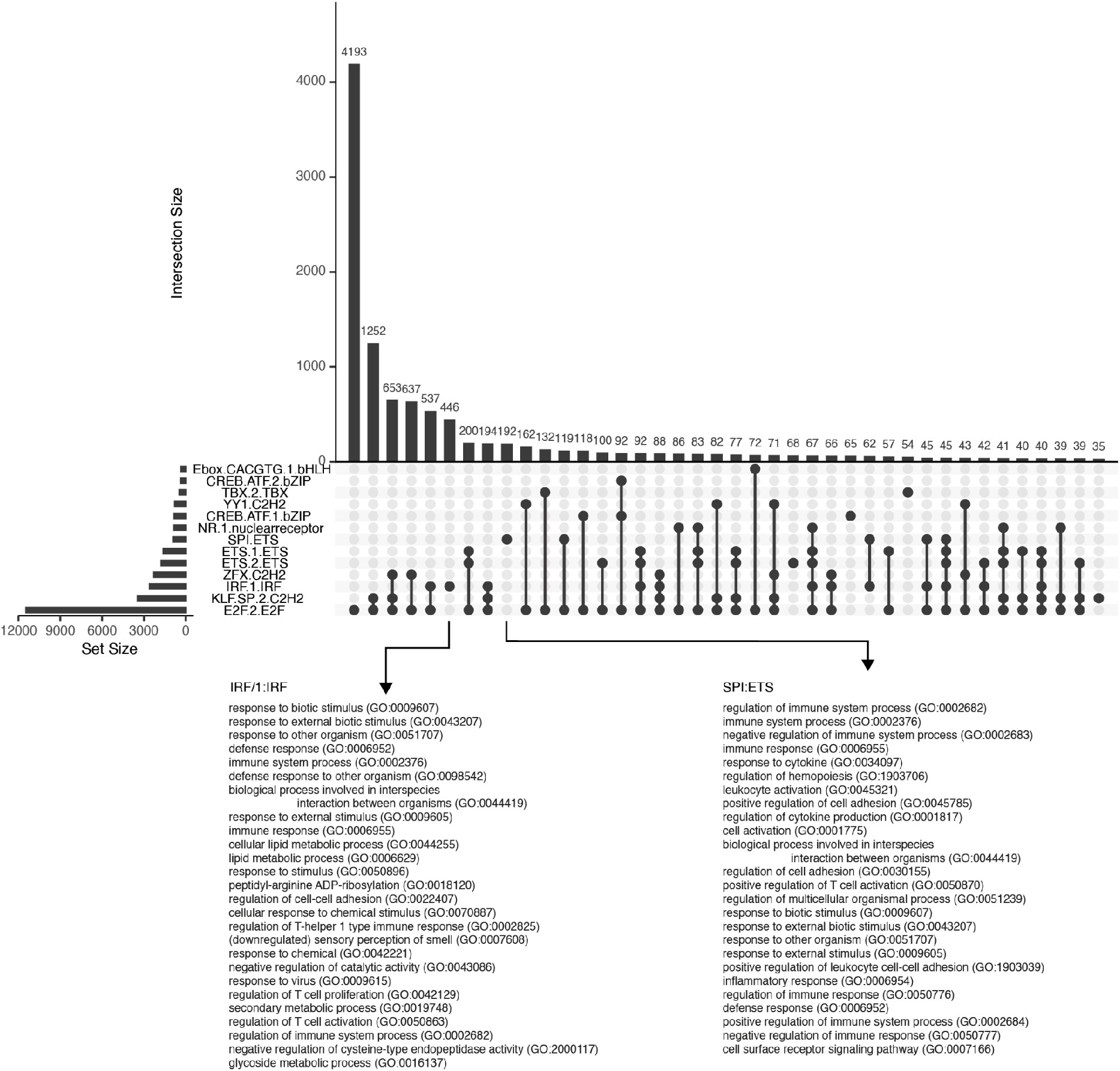
Co-occurring motif families in mouse promoters (Methods). GO-term biological process enrichment for the two given gene sets are shown below, ordered for FDR-adjusted *p*-values. All GO-terms are enriched except for ‘sensory perception of smell’ for IRF/1:IRF (this one is underrepresented).

**Figure S8:**
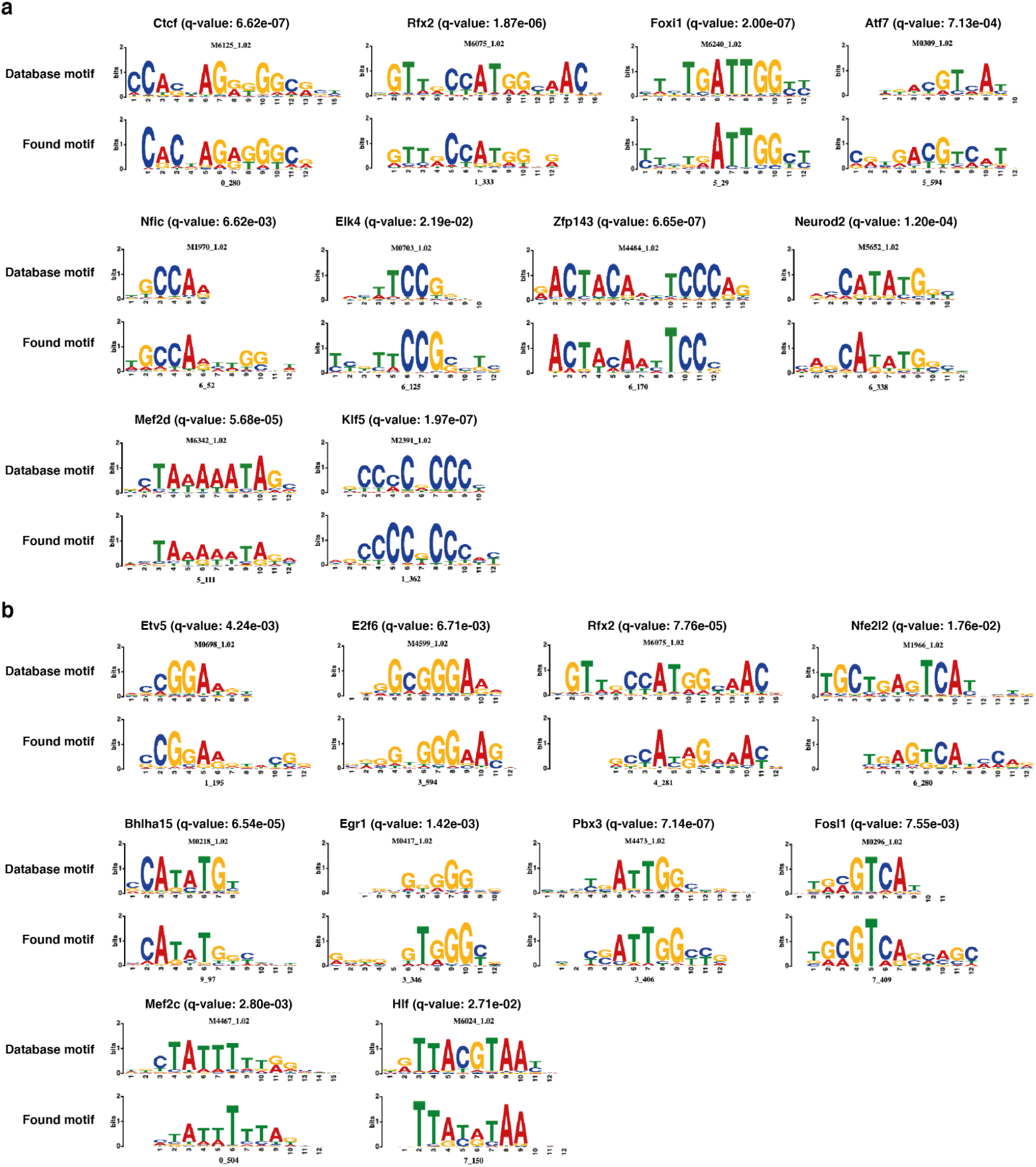
Example motif alignments for the SNARE-seq (*a*) P0 and (*b*) adult datasets.

**Figure S9:**
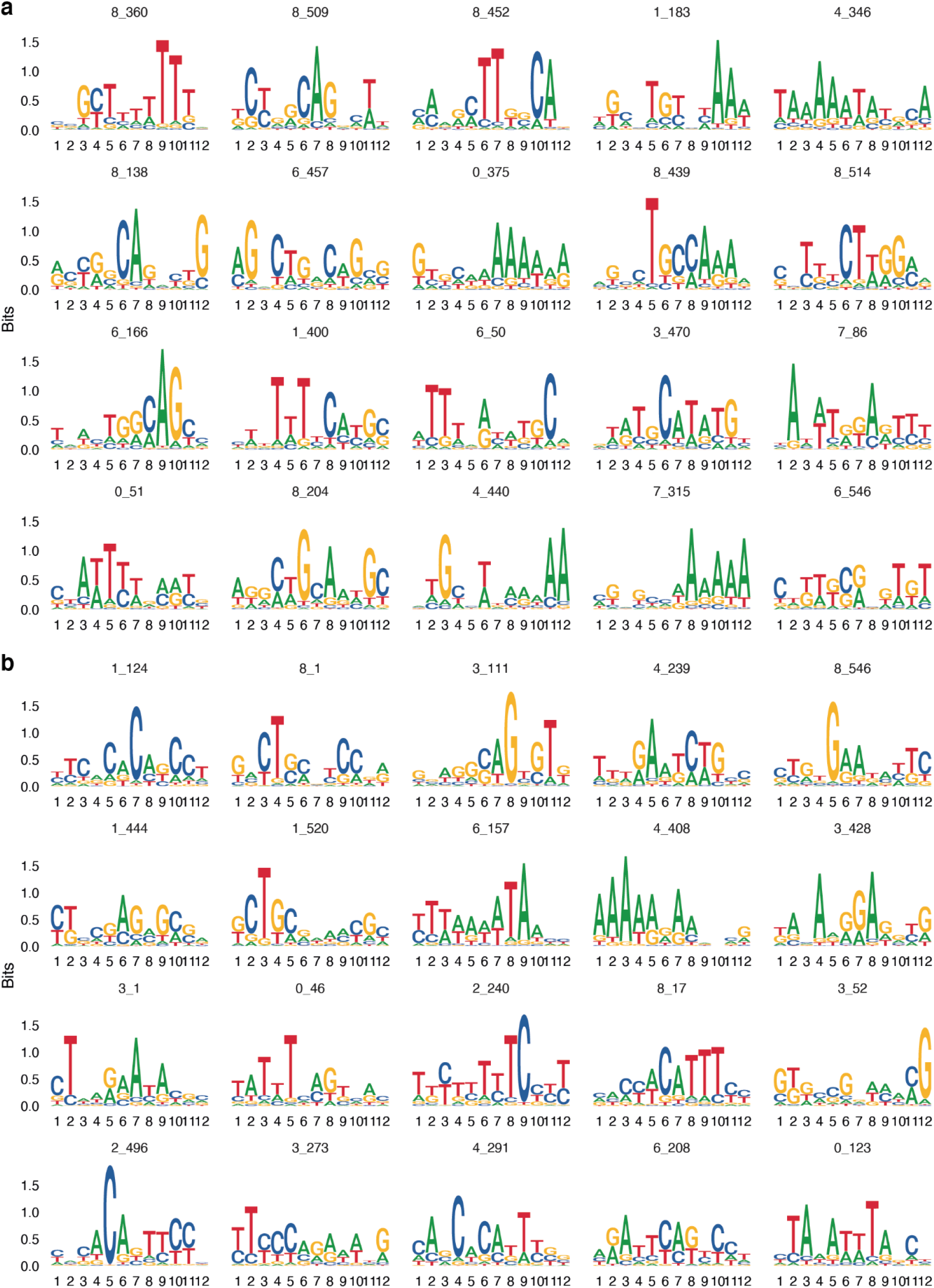
Examples of randomly selected motifs with high influence scores that did not align to CIS-BP for the (*a*) P0 dataset and (*b*) adult dataset from SNARE-seq. Partial motif matches are found: for example, in P0, motif 4_346 is a partial match to MEF2 family motifs, and motif 3_470 has an E-box motif (CANNTG). For the adult dataset, 0_46 is a partial match to MEF2 family motifs.

**Figure S10:**
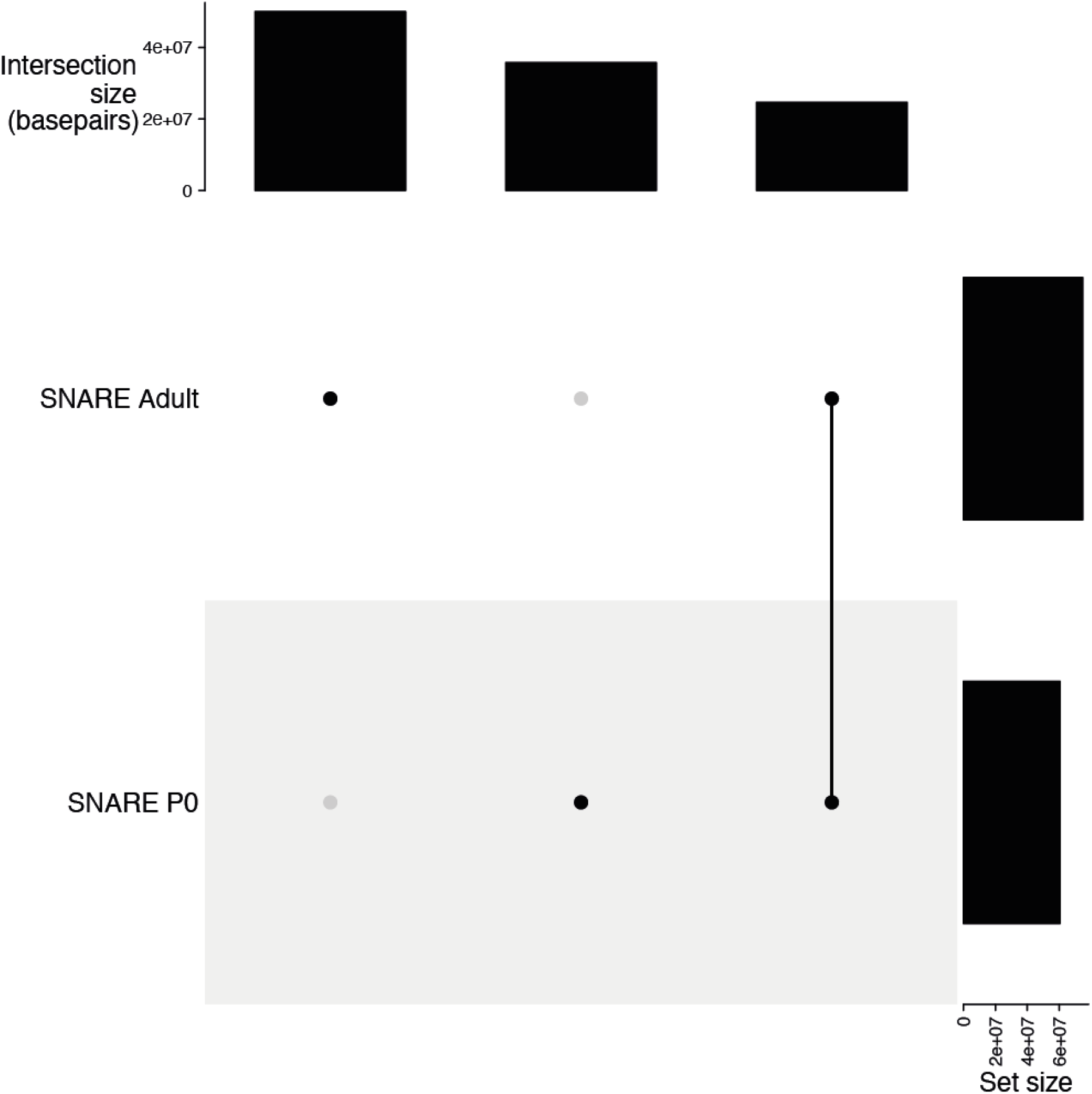
Overlap of open chromatin regions from different time points. Set sizes are calculated as the total number of base pairs overlapping in the two datasets.

**Figure S11:**
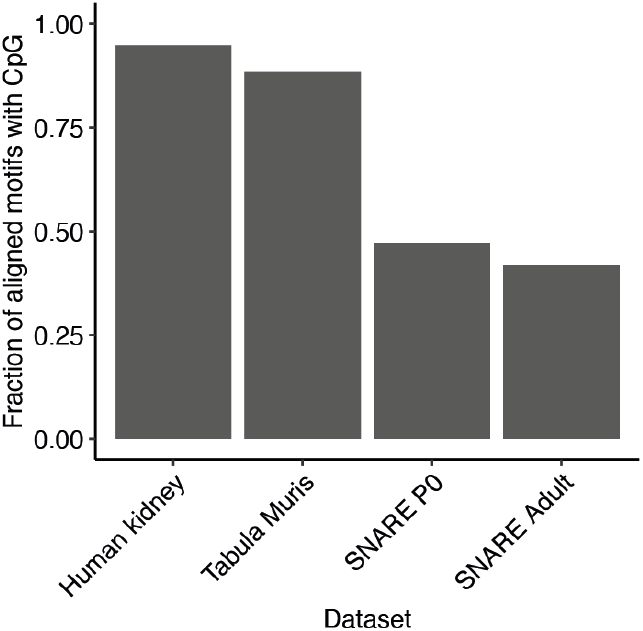
CpG count for the motifs found using the different datasets.

**Figure S12:**
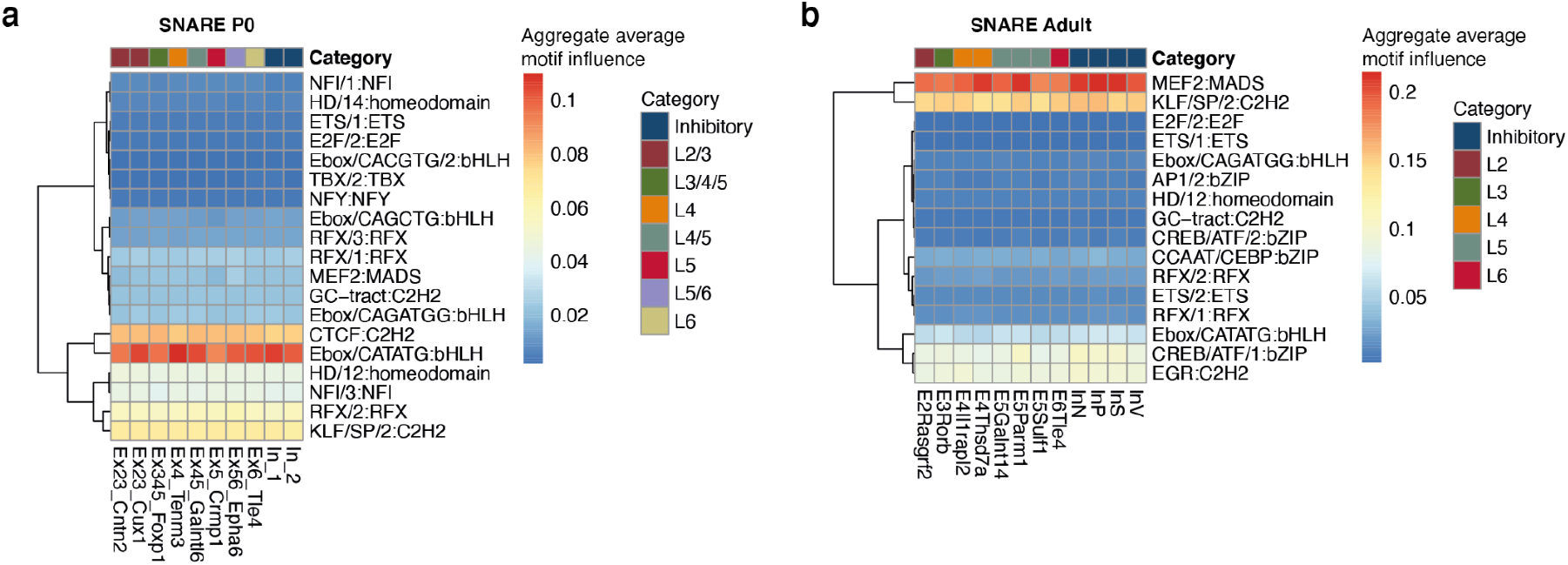
(*a*) Aggregate motif weights of motif clusters averaged in cell types for the SNARE-seq P0 dataset. (*b*) Aggregate motif weights of motif clusters averaged in cell types for the SNARE-seq Adult dataset.

## Supplementary tables

**Table S1:** Assignment of cell types to categories for the human kidney dataset

| **Cell type** | **Category** |
| --- | --- |
| Ascending vasa recta endothelium | Endothelium |
| B cell | Immune |
| CD4 T cell | Immune |
| CD8 T cell | Immune |
| cDC2 | Immune |
| CNT/PC - proximal UB | Nephron epithelium |
| Cap mesenchyme | Nephron progenitor |
| Connecting tubule | Nephron epithelium |
| Descending vasa recta endothelium | Endothelium |
| Distal S shaped body | Nephron progenitor |
| Distal renal vesicle | Nephron progenitor |
| Endothelium | Endothelium |
| Epithelial progenitor cell | Nephron progenitor |
| Fibroblast 1 | Stroma |
| Fibroblast 2 | Stroma |
| Glomerular endothelium | Endothelium |
| Indistinct intercalated cell | Nephron epithelium |
| Innate like lymphocyte | Immune |
| Loop of Henle | Nephron epithelium |
| MNP-a/classical monocyte derived | Immune |
| MNP-b/non-classical monocyte derived | Immune |
| MNP-c/dendritic cell | Immune |
| MNP-d/Tissue macrophage | Immune |
| Macrophage 1 | Immune |
| Macrophage 2 | Immune |
| Mast cells | Immune |
| Medial S shaped body | Nephron progenitor |
| Megakaryocyte | Immune |
| Monocyte | Immune |
| Myofibroblast | Stroma |
| Myofibroblast 1 | Stroma |
| Myofibroblast 2 | Stroma |
| NK cell | Immune |
| NKT cell | Immune |
| Neutrophil | Immune |
| pDC | Immune |
| Pelvic epithelium | Nephron epithelium |
| Pelvic epithelium - distal UB | Nephron progenitor |
| Peritubular capillary endothelium 1 | Endothelium |
| Peritubular capillary endothelium 2 | Endothelium |
| Podocyte | Nephron epithelium |
| Principal cell | Nephron epithelium |
| Proliferating B cell | Immune |
| Proliferating NK cell | Immune |
| Proliferating cDC2 | Immune |
| Proliferating cap mesenchyme | Nephron progenitor |
| Proliferating distal renal vesicle | Nephron progenitor |
| Proliferating fibroblast | Stroma |
| Proliferating macrophage | Immune |
| Proliferating monocyte | Immune |
| Proliferating myofibroblast | Stroma |
| Proliferating stroma progenitor | Stroma |
| Proximal S shaped body | Nephron progenitor |
| Proximal UB | Nephron progenitor |
| Proximal renal vesicle | Nephron progenitor |
| Proximal tubule | Nephron epithelium |
| Stroma progenitor | Stroma |
| Thick ascending limb of Loop of Henle | Nephron epithelium |
| Type A intercalated cell | Nephron epithelium |
| Type B intercalated cell | Nephron epithelium |

**Table S2:** Assignment of cell types to categories for the Tabula Muris dataset

| **Cell type** | **Category** |
| --- | --- |
| bladder basal cell of urothelium | epithelial |
| bladder bladder cell | epithelial |
| bladder mesenchymal cell | connective |
| brainmicroglia microglial cell | immune |
| brainneurons astrocyte of the cerebral cortex | macroglial |
| brainneurons brain pericyte | pericyte |
| brainneurons endothelial cell | endothelial |
| brainneurons neuron | neuronal |
| brainneurons oligodendrocyte | macroglial |
| brainneurons oligodendrocyte precursor cell | macroglial |
| colon enterocyte of epithelium of large intestine | epithelial |
| colon epithelial cell of large intestine | epithelial |
| colon large intestine goblet cell | epithelial |
| fat B cell | immune |
| fat endothelial cell | endothelial |
| fat granulocyte | immune |
| fat mesenchymal stem cell of adipose | connective |
| fat myeloid cell | immune |
| fat natural killer cell | immune |
| fat neutrophil | immune |
| fat T cell | immune |
| heart cardiac muscle cell | muscle |
| heart endocardial cell | endothelial |
| heart endothelial cell | endothelial |
| heart epicardial adipocyte | adipose |
| heart fibroblast | connective |
| heart leukocyte | immune |
| heart smooth muscle cell | muscle |
| kidney kidney tubule cell | epithelial |
| liver endothelial cell of hepatic sinusoid | endothelial |
| liver hepatocyte | epithelial |
| lung endothelial cell | endothelial |
| lung stromal cell | connective |
| lung type II pneumocyte | epithelial |
| mammary basal cell | epithelial |
| mammary luminal epithelial cell of mammary gland | epithelial |
| mammary stromal cell | connective |
| marrow B cell | immune |
| marrow Fraction A pre-pro B cell | immune |
| marrow granulocyte | immune |
| marrow hematopoietic stem cell | immune |
| marrow monocyte | immune |
| marrow natural killer cell | immune |
| marrow neutrophil | immune |
| marrow T cell | immune |
| muscle B cell | immune |
| muscle endothelial cell | endothelial |
| muscle mesenchymal stem cell | connective |
| muscle skeletal muscle satellite cell | muscle |
| muscle skeletal muscle satellite stem cell | muscle |
| pancreas pancreatic A cell | endocrine |
| pancreas pancreatic acinar cell | exocrine |
| pancreas pancreatic D cell | endocrine |
| pancreas pancreatic ductal cell | epithelial |
| pancreas pancreatic PP cell | endocrine |
| pancreas type B pancreatic cell | endocrine |
| skin basal cell of epidermis | epithelial |
| skin epidermal cell | epithelial |
| skin keratinocyte stem cell | epithelial |
| spleen B cell | immune |
| spleen T cell | immune |
| thymus T cell | immune |
| tongue basal cell of epidermis | epithelial |
| tongue keratinocyte | epithelial |
| trachea epithelial cell | epithelial |
| trachea leukocyte | immune |
| trachea stromal cell | connective |

